# A Fully Endovascular Neural Interface

**DOI:** 10.64898/2026.06.07.730604

**Authors:** John W. Stanton, Giovanni Talei-Franzesi, Eleonora F. Spinazzi, Jennifer Haupt, Darren Orbach, Jonathan Sisti, Morgan Jamiel, Emily Zhang, Alexander Romanov, Sean D. Lavine, E Sander Connolly, Elisa Konofagou, Edward S. Boyden, Kenneth L. Shepard

## Abstract

Electrical stimulation of neural circuits is expanding therapeutic strategies to modulate brain, autonomic, and immune functions. Devices delivered endovascularly offer a less invasive alternative to conventional implanted electrodes, while delivering spatio-temporal specificity superior to noninvasive techniques. We demonstrate a fully endovascular sub-1-mm^3^ implant, utilizing ultrasound for wireless power delivery and data telemetry in a fashion invariant to device orientation. The implant consists of piezoelectric transducers, an energy storage capacitor, an application-specific integrated circuit, and electrodes packaged on a 7-µm-thick polyimide scaffold. The implant can be delivered through a microcatheter in a manner analogous to conventional neurovascular stents, and self-expands upon deployment to appose the vessel walls. We demonstrate intravascular stimulation of the autonomic nervous system from the carotid artery, achieving modulation of blood pressure in rabbits. This approach establishes a broadly applicable platform for neural interfaces enabling both stimulation and recording.

## INTRODUCTION

The autonomic nervous system (ANS) plays a vital role in the maintenance of homeostasis by regulating critical organ and tissue functions and is consequently involved in the pathogenesis of many cardiovascular, inflammatory, and metabolic disease processes ^1,2^. For example, the role of the ANS in regulating cardiovascular output, including heart rate, blood pressure, and vascular tone, directly implicates the ANS in arterial hypertension (aHT), or elevated blood pressure which affects roughly half of the adult population in the United States ^3^. Despite notable advances in the pharmacologic treatment of aHT, a substantial number of patients fail to achieve blood pressure (BP) control on three or more antihypertensive drugs^4,5^ Device-based approaches based on peripheral nervous system (PNS) neuromodulation, known as electroceuticals, can provide an alternative to such pharmacologically-resistant conditions with easily identifiable nervous system targets if these devices can be made less invasive in their deployment.

Neurovascular bundles bind nerves and vascular structures together such that they travel in tandem; for this reason, the vasculature provides an access pathway to nervous system tissue throughout the body^6^. Vascular neural interfaces have the advantage of having more chronically stable impedances compared with direct electrode contact because no nervous tissue is disturbed or irritated in the process of implantation ^7^. Acute transvascular neural recording and stimulation has been demonstrated in both animals and humans for both central-nervous-system (CNS) and PNS targets, with recent work suggesting that single-unit decoding from the microvasculature is feasible ^8^. Recently, a sixteen-electrode intravascular neural interface designed for long-term implantation was demonstrated for the first time^9,10^; this system places electrodes inside the vessel while the interfacing electronics are housed entirely extravascularly in a canister implanted subcutaneously in the chest. Such devices employ tethered electrode interconnects, requiring wire bundles to be drawn through the vasculature from the implant for measurement. These long and necessarily thin wires present a permanent risk of thrombosis and ischemic infarction for implants placed on the arterial side and of venous stroke and hemorrhage for those on the venous side. The risks associated with these procedures in addition to the tissue displacement of the electronics canister severely limit the claims of minimal invasiveness, and the risk of mechanical fatigue and breakage over device lifetime introduces an additional failure mode ^11^. Delivery to small diameter vessels is particularly challenging with this approach, restricting the potential deployment endpoints to major vascular pathways. These devices also face scalability challenges, as channel counts are fundamentally constrained by the ability to route and escape an increasing number of wires from the vasculature without occlusion of blood flow or significant increase in the risk of hemodynamic complications.

By taking a volumetrically efficient approach to the bioelectronic implant design, however, we can remove traditionally large volume components and miniaturize the wireless system architecture such that it fits entirely within the vasculature and can be inserted percutaneously without the need for additional implanted hardware or surgical complexity. On-board batteries and their hermetically sealed housing comprise the largest contribution to the volume of traditional implantable devices. Maximizing volumetric efficiency requires wirelessly powering a fully-integrated system with optimized thin-film biocompatible encapsulation and interfaces.

Advances in wearable ultrasound (US) devices enable acoustic power and data transmission to be effectively employed for implantable devices, which delivers several advantages over more traditional electromagnetic (EM) approaches ^12^. Because of the large difference in phase velocity between US and EM energy, US has a wavelength of only ∼0.77mm at 2 MHz compared with 25 mm for 2-GHz EM radiation ^13^. Efficient coupling to far-field energy is dimensionally limited by wavelength. US supports far-field energy coupling at MHz carrier frequencies to millimeter-length-scale transducer materials, providing for superior power transfer efficiency and high-signal-to-noise-ratio bidirectional communication with signal levels limited only by attenuation in tissue. Furthermore, the US Food and Drug Administration (FDA) limitations on power flux are also substantially higher for acoustic energy compared to EM energy (7.2 mW/mm^2^ for US compared with 0.1 mW/mm^2^ for EM), while having substantially lower attenuation through soft tissue (2 dB/cm for a 1-mm-sized implant with US compared with 14.6 dB/cm for a 10-mm-sized implant operating at 3 GHz with EM ^14^). By employing US, we are able to safely deliver sufficient power to achieve target stimulation parameters at volumes compatible with chronic implantation within small-diameter vessel structures.

Clinical translation of an endovascular technology requires that the implant must be deliverable with small-diameter catheters using conventional techniques, while maintaining a minimal profile to avoid the risk of acute and chronic hemodynamic and thromboembolic complications. Modern cardiovascular stents used in percutaneous coronary intervention typically have strut thicknesses of 60-80 µm and target vessels are in the 2-4.5 mm diameter range, and while the specific implications on hemodynamics are assessed on a per-implant basis, the cross-sectional area presented inside the vessel lumen serves as a starting basis to predict the viability of the implant. For conventional stenting technologies, values from 0.023 - 0.18 mm^2^ ^15^ can be taken as typical. The only other wireless stimulator device reported for endovascular applications to-date ^16^ has a cross sectional area of 6.45 mm^2^ (3-mm by 2.15-mm), making it substantially too large for safe chronic deployment within most vascular delivery targets and consequently was not demonstrated as a fully endovascular deployment.

In this study, we introduce the first fully endovascular implant without any escaping wires, electrodes, or large volume structures outside of the vessel, while maintaining a small enough cross-section to be a promising candidate for chronic clinical applications. Used as a vascular neural interface (VNI), this implantable stent-like device is designed to be viable in either the CNS or PNS and to be delivered into vessels as small as ∼1.4-mm in diameter. The VNI device is comprised of an array of three acoustic transducers, allowing for angle-insensitive powering and telemetry, and a custom application-specific integrated circuit (ASIC). The overall system package is less than 1 mm^3^ in volume, weighs 6.8 mg, and has a cross-sectional profile corresponding, at its maximum, to ∼5.3% of the area of a 2-mm-diameter lumen vessel (∼.14 mm^2^ of cross section), less than the profile of ultra-thin strut stents in clinical use (∼11.6% for 60 µm-thick struts completely encircling the lumen ^15^). In a rabbit model system, we show the modulation of blood pressure through electrical stimulation of the ANS from the VNI adjacent to its nerve target.

## RESULTS

The VNI system is a vascularly deployed nervous system interface that is wirelessly powered through US and controlled by a remote acoustic transducer. An overview of the VNI system and its implementation are shown in **Fig. 1a**. The external US system establishes an acoustic link with the implanted VNI device which powers the implant, provides a bidirectional communication link with the VNI implant, and controls its stimulation output, as shown in **Fig. 1b**. The implant, shown in **Fig. 1c**, consists of three identical lead-magnesium niobate-lead-titanate (PMN-PT) piezoceramic transducers (see **Supplementary Discussion S1** and **S2**) designed to deploy in a spatially offset manner around the vessel circumference (see **Supplementary Discussion S3**); a custom designed ASIC to generate a 5-V stimulation supply and to provide stimulation pulses with controllable parameters (see **Supplementary Discussion S4**); a pair of gold metallic electrodes, analogous to ring electrodes used for deep-brain-stimulation (DBS), which span the circumference of the vessel (see **Supplementary Discussion S5**); and an energy storage capacitor to decouple the stimulation supply. The gold stimulation electrodes are 1.5-mm wide on a pitch of 3.7-mm. A custom fabricated mechanically flexible thin-film polyimide substrate integrates the ASIC with the external components via a single layer of gold interconnects. The substrate is highly flexible and conforms to whatever surface on which it is placed when laid flat, as seen in **Fig. 1d**. However, the intrinsic stresses in the polyimide film when rolled along its lengthwise axis (see **Supplementary Discussion S6**) allow the VNI implant to unfurl into tight apposition with the vessel wall when released from a microcatheter delivery vehicle, as shown in **Fig. 1e**.

**Figure 1.**
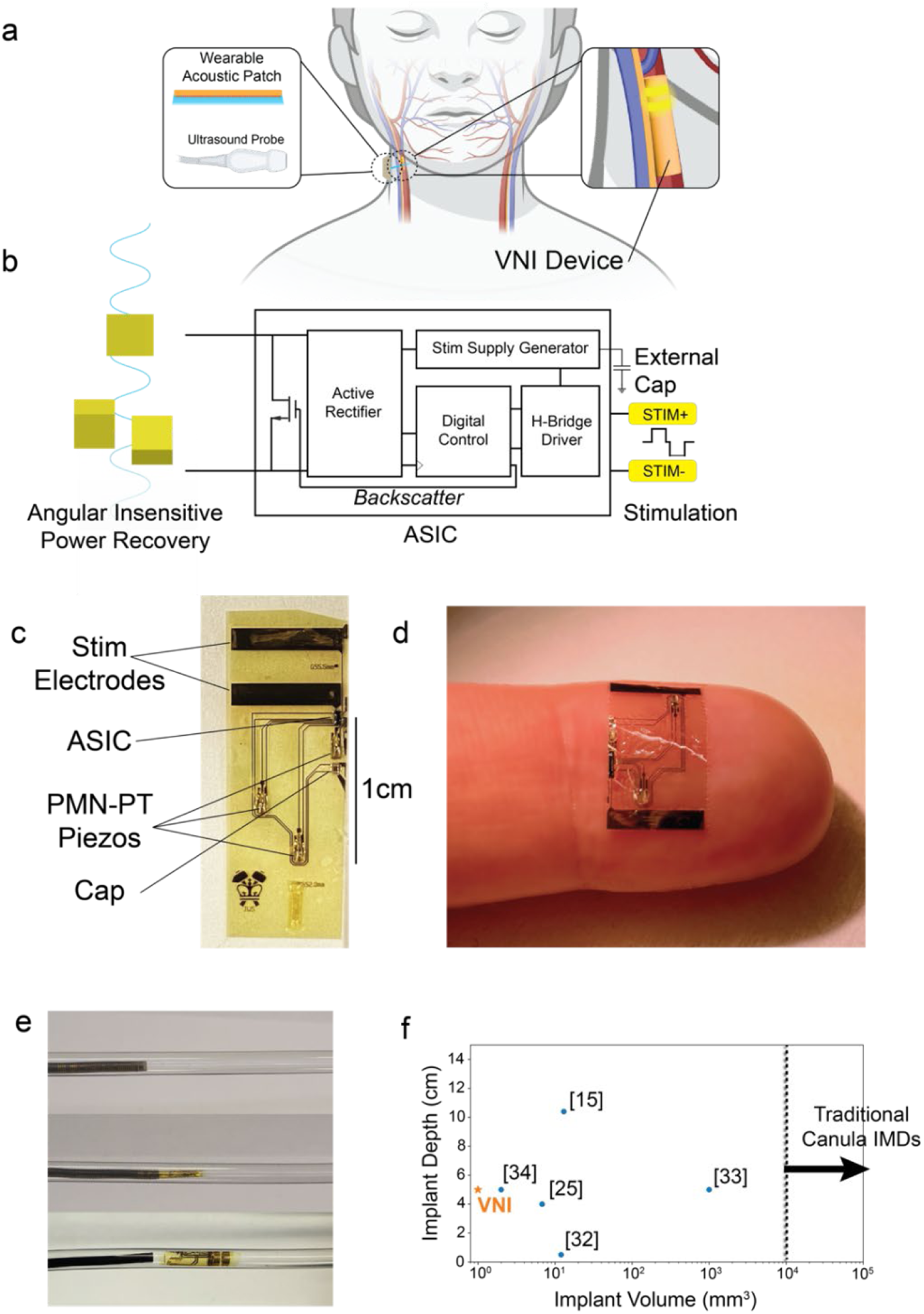
Design of the VNI system. **a)** Diagram depicting the VNI system, placed in an artery with an external acoustic transducer powering and communicating with the implanted VNI. This transducer could take the form of a conventional ultrasound probe (as employed in the studies here) or emerging wearable ultrasound “patch” devices. Anatomical rendering is created in-part with BioRender.com **b)** System design of the VNI implant developed here which includes an ASIC packaged together with piezoelectric transducers, electrodes, and an energy-storage capacitor. **c)** Layout of the components of the VNI system in the vessel, with structural components inside the vascular structure and the electrodes facing the vessel wall. **d)** Demonstration of flexibility of VNI device curled around finger for scale. **e)** Delivery of VNI into 2-mm-inner-diameter tubing. **f)** Comparison of the VNI system to other implanted US-powered stimulators ^16,18,38–40^ using the metrics of reported maximum implanted depth and implant volume.

In this study, the VNI device is positioned and delivered within the common carotid artery at locations corresponding to cervical levels C6-C3 using vascular navigation facilitated by fluoroscopic guidance starting from a femoral-artery access point. Conventional vascular navigation techniques are used to direct the VNI delivery system to the target endpoint and to release the VNI implant in a manner similar to self-expanding stents for clinical use, allowing it to mechanically brace against the vessel walls. Once delivered, a fluoroscopic contrast agent is injected to check the vessel for patency. While this study deploys the VNI device in an artery, venous deployments are also possible and might present a lower-risk pathway for clinical translation. The improvement in volumetric efficiency over existing implantable acoustically powered stimulation devices enable fully endovascular delivery to a vast range of target sites, and the potential for safe chronic usage in a clinical setting (**Fig. 1f**).

### Acoustic Protocol

Once deployed, the VNI interfaces with an external ultrasound system, which in this study is a commercial ultrasound probe, coupled to the skin with acoustic coupling gel, controlled by a Verasonics programmable ultrasound system. This research-grade US system provides per-element beam-forming, pulse-echo reception, digitization, and image reconstruction, shown in **Fig. 2a**. In addition to powering the implant, the external ultrasound system enables a bidirectional data link. A timing diagram of the acoustic protocol is shown in **Fig. 2b**. To program the implant, a pulse-width-modulated (PWM) downlink pulse sequence at a 2-MHz carrier is used in which acoustic pulses longer than 16 cycles encode logic ‘1’, while pulse lengths shorter than 16 cycles encode logic ‘0’; there is a 6-μs interval between pulses. An example pulse sequence is shown in **Fig. 2b**. The probe also receives uplink data from the VNI device by receiving backscattered signals from the implant; this backscatter is produced by load-shift-keying (LSK), modulating the VNI transducer’s acoustic impedance by modulating its electrical impedance with a shorting transistor in the acoustic front-end circuits of the ASIC.

**Figure 2.**
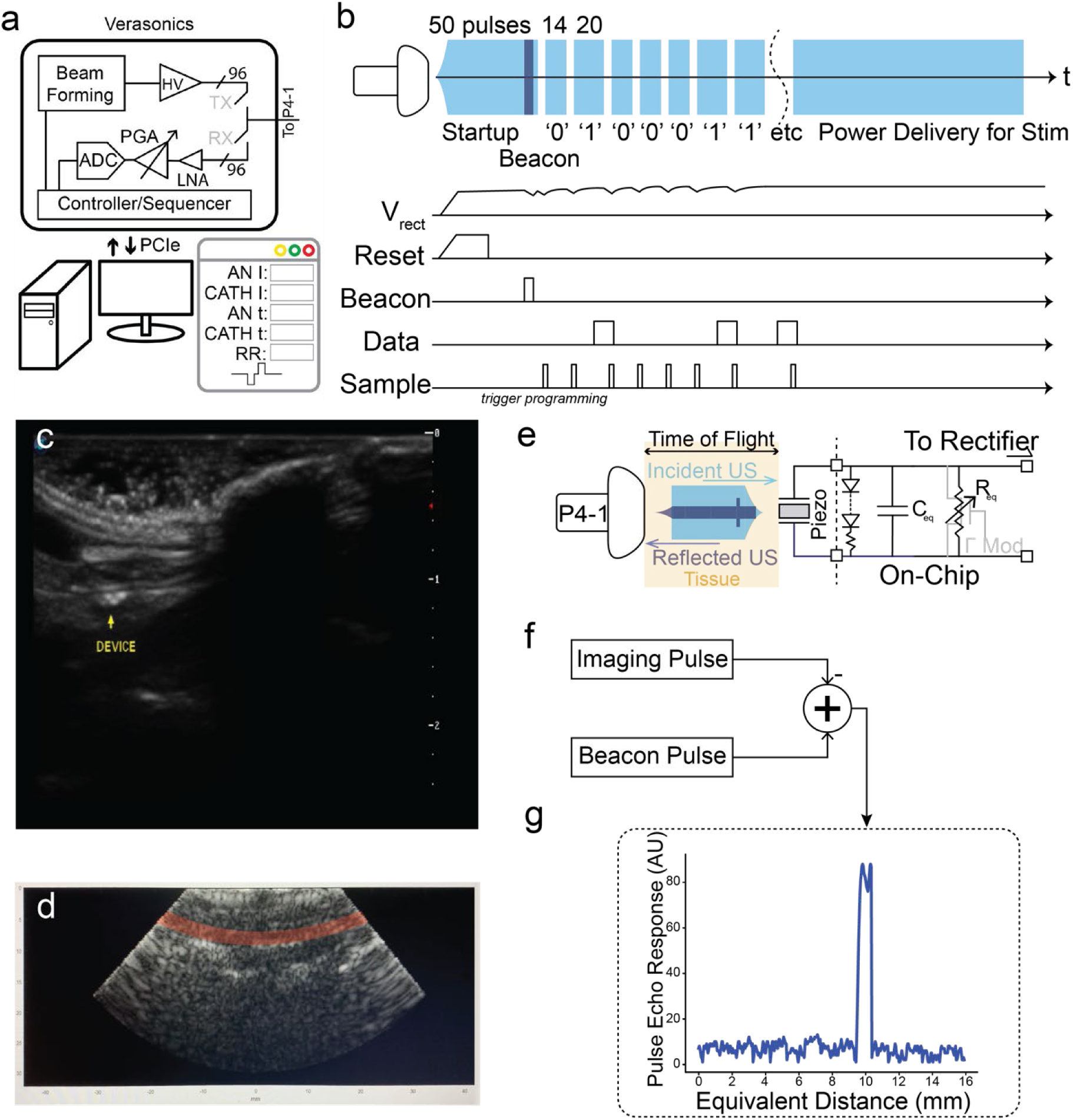
Acoustic protocol for the VNI system. **a)** Functional block diagram of the external ultrasound system and custom control user interface. **b)** Timing diagram of the acoustic pulses, showing the interaction between the incident ultrasound. **c)** 10-MHz high-resolution B-Mode image of the VNI device. **d)** 2-MHz low-resolution B-Mode image of the vasculature of the NZWR, with the common carotid highlighted. **e)** Functional image of the beacon pulse system **f)** Pulse-echo-response processing approach to detect beacon LSK response. **g)** Representative pulse-echo response from the VNI implant during device-discovery phase plotted

A high-frequency (10 MHz) medical-grade acoustic imaging system is able to resolve the VNI device (**Fig. 2c**). However, 2-MHz imaging (**Fig. 2d**) has insufficient resolution to identify any structural elements in the VNI since all echogenic components of the VNI are on the order of a single acoustic wavelength at this frequency. Due to the pulsatility of the carotid artery, we are able to resolve the ∼2-mm carotid artery itself in a B-Mode imaging mode at 2 MHz. To make the implant more visible at this frequency, after initial power-up, the VNI device sends a “beacon” response in which the backscatter is enhanced for approximately 8-µs (**Fig. 2e**). The general procedure for locating the device with ultrasound begins with a two-pulse plane-wave imaging sequence, in which the entire probe aperture emits an imaging pulse to record the baseline pulse-echo response of the tissue bed. Then, a fifty-pulse plane-wave sequence is emitted by the acoustic probe, which activates the implant and provides sufficient clock cycles to trigger the beacon response pulse. The pulse-echo-response data is recovered across the transducer aperture, and the difference between the baseline and the beacon pulse sequence is calculated (**Fig. 2f**). The resulting volumetric field data is analyzed for the signature of the implant, with each transducer element’s raw response difference data analyzed with thresholding to search for a signal from the implant. Each element’s position along the transducer aperture and pulse echo response time-of-flight (determined by the Verasonics sampling rate and acoustic phase velocity) can be used to determine the implant’s depth and translational position. If no beacon response is found, the probe is repositioned on the neck, either by translating its position or by adjusting its rotational orientation.

When the VNI is within the transducer’s field of view, the implant has a typical response signal-to-noise ratio (SNR) of 15.6 dB. A representative difference trace, and corresponding pulse-echo response, is shown in **Fig. 2g**. Typically, we aim to localize the VNI device under the center of the acoustic probe, moving the probe if the beacon response is detected from end-elements. After determining the VNI-device’s location using the beacon-mode response, the acoustic probe is typically immobilized with a stand or clamp. However, there are many possible ray-lines between the external probe and the implant, and we have been able to maintain the VNI-implant interface while simply holding the acoustic probe by hand.

The VNI platform has two functional modes. The first mode is a continuous acoustic excitation mode, in which the ultrasound probe programs the implant and then provides continuous power while the implant stimulates according to a programmed excitation waveform. The second mode is the pulse-excitation mode, in which the implant is programmed for each delivered US excitation sequence, and an electrical stimulation pulse is delivered. This minimizes heating at the probe/tissue interface for low-repetition-rate stimulation patterns (see **Supplementary Discussion S7**).

### Functioning of the CMOS ASIC

A block diagram of the VNI ASIC is shown in **Fig. 3a**. Incident ultrasound is harvested by the piezoelectric transducers and converted to an AC input voltage at the carrier frequency (*f_carrier_*) of 2 MHz, which is used to generate an unregulated powering voltage of approximately 1.5-V through an active rectifier with diode-connected PMOS devices to ensure startup from a cold state. The buffered comparator in each active rectifier clocks a D-flip-flop to also generate a half-rate 50% duty cycle system clock at *f_carrier_*/2 with a clock period of 1 µs. The powering voltage feeds a 0.5-V voltage reference and a low-dropout (LDO) regulator to generate a 1.2-V logic supply.

**Figure 3.**
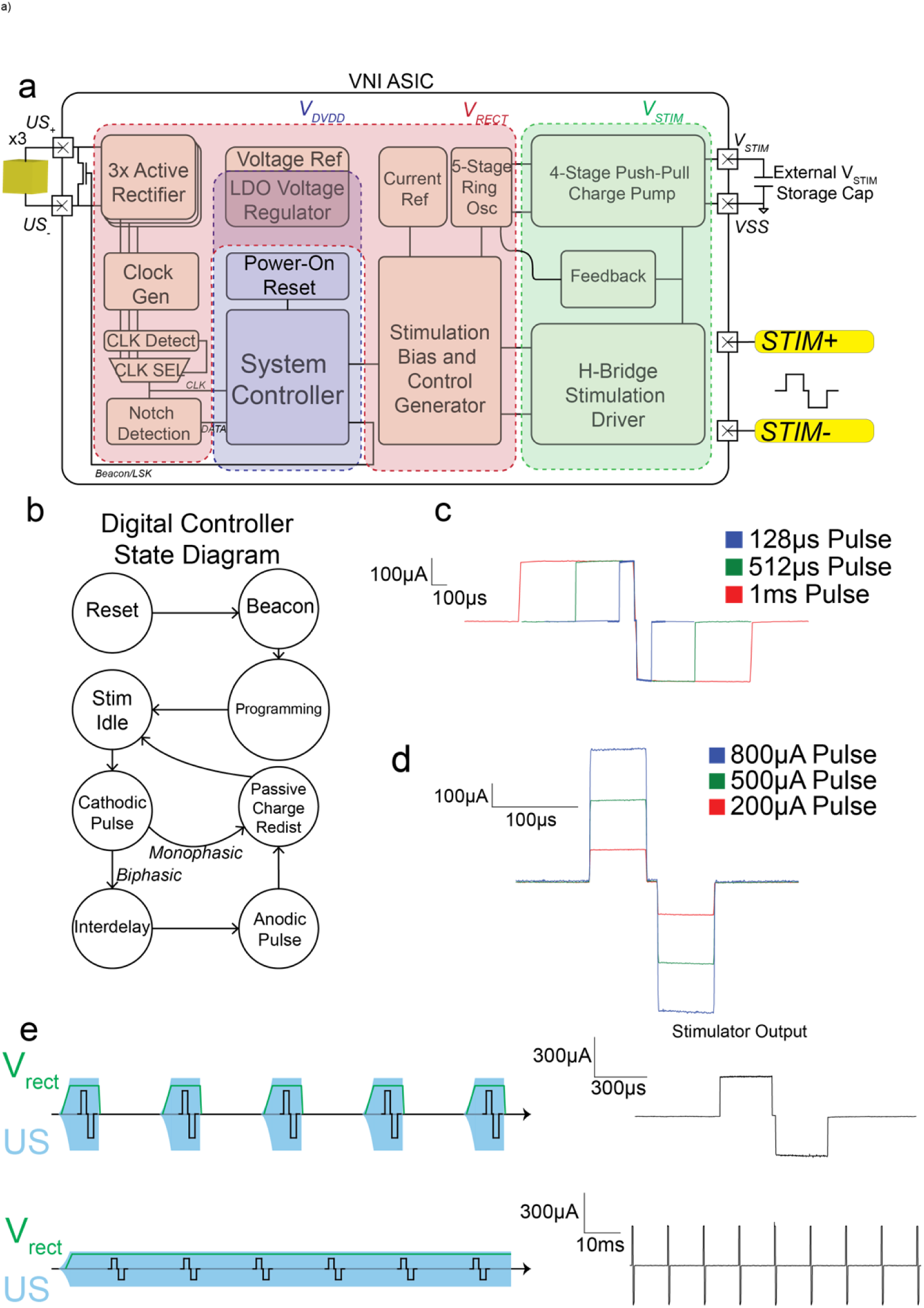
Design of the VNI endovascular implant. **a)** Block diagram of the VNI ASIC. **b)** State transition diagram for the VNI. **c)** Symmetric stimulator output at different programmable timings. **d)** Symmetric stimulator output at different programmable output currents **e)** Output of the VNI implant in *in-vitro* testing of both operational modes. Top: pulsed mode, in which a single stimulation pulse is delivered with each ultrasound pulse. Bottom: continuous mode in which pulse delivery is programmed into the VNI device and delivered with continuous ultrasound stimulation.

An additional 5-V supply for the stimulator is generated from a four-stage push-pull charge pump that is clocked by a current-starved ring oscillator. The oscillator is designed to increase in frequency to compensate for droops in the stimulation supply (e. g., upon initial startup or as the result of a stimulation event). As the ring oscillator increases in frequency, more charge is delivered to the stimulation supply. However, because the charge pump can operate at frequencies faster than the rate at which the acoustic front end can replenish the rectified system voltage, the charge pump could brown-out the rest of the ASIC. An under-voltage supply monitor is used to throttle the charge-pump oscillator frequency if the charge-pump is pulling too much charge away from the system supply. To save power, the stimulator biasing is off when not in use. The supply monitor also ensures that the stimulation supply is at least 3.8-V before stimulation is permitted; no pulse is delivered until charge-balancing can be guaranteed. The ASIC consumes 294 µW of power while the charge pump is active.

During VNI programming, an analog watchdog timer detects the 6-μs interval between bits. Programming is initiated at the first of these intervals. After programming, the stimulator delivers stimulation pulses based on the decoded settings until the system powers down. A state diagram of the ASIC digital controller that governs the behavior of the implant is shown in **Fig. 3b**. If the VNI is configured in continuous stimulation mode, the stimulator will provide a digitally programmable biphasic constant-current pulse sequence with amplitude between 100 μA and 800 μA in steps of 100 μA and pulse widths between 50-µs to 1-ms for the anodic and cathodic phases at a programmable repetition rate (100 Hz to 1 kHz). If the VNI is configured in pulse excitation mode, the same biphasic pulse parameters can be delivered to the tissue; however, the maximum pulse repetition rate is 154 Hz. The two different modes of operation are illustrated in **Fig. 3e**.

The stimulator is biased from a 100-nA on-implant-generated current reference, which is multiplied by tunable current mirrors. Various stimulator output pulse widths can be delivered, as shown in **Fig. 3c**, along with different stimulator pulse amplitudes, as shown in **Fig. 3d**. After stimulation is delivered the ASIC weakly shorts the stimulator anode and cathode for passive charge balancing. More details on the circuit architecture are provided in **Supplementary Discussion S4**.

### Rotationally Invariant Design

Ultrasound is an emerging approach for communicating with and powering implantable biomedical systems ^17–19^; however, the devices described to-date have generally had, and required, the advantage of being oriented for optimal acoustic power delivery and telemetry during surgical implantation. While such careful alignment is possible in an acute research setting, it may complicate translation to chronic clinical use. Potential alignment issues are exacerbated in the catheter-based deployment of the VNI because the delivery vehicle generally affords little control of the rotational orientation of the VNI, and once unsheathed into the vessel, the unfurling of the device inside the vessel can occur randomly. The VNI device is designed for rotationally-invariant powering, an approach which may also prove beneficial for other kinds of ultrasound-powered medical implants. To accomplish this, three piezeoelectric transducers are integrated on the VNI package and radially offset by 120 degrees from each other around the circumference of the expanded device. The transducers are also offset along the length of the polymeric stent in order to minimize the cross-sectional profile at any discrete point along the implant’s length (see **Supplementary Discussion S3**), reducing the risk of thrombosis. The system clock is derived from the incident ultrasound carrier frequency. An activity detection circuit identifies which piezoelectric transducer and active rectifier are receiving power, and selectively multiplexes a single clock from the buffered active-rectifier comparator such that system clocking remains glitch-free. While the clock is generated from the dominant rectifier, energy can be continuously harvested from all three transducers.

### Flexible Polymeric Package Design

A microfabrication process flow diagram detailing the steps required to fabricate the thin-film polymeric stent is provided in **Supplementary Fig. S9**. The volumetrically significant components of the system (the ASIC chip, the piezoelectric transducers, and the energy storage capacitor) are all internal to the vessel while the electrodes, which span the circumference of the vessel, face the vessel wall as shown in **Fig. 1c**. In order to minimize hemodynamic disruption and reduce the risk of acute or chronic complications, the chip, piezoelectric transducers, and capacitor are offset along the length of the VNI such that only one occlusive component obstructs vascular flow at any cross-section along the length of the implant.

To further reduce implant volumes, interconnection of components to the package relies on the use of anisotropic conductive films (ACFs). The ASIC is flip-chip bonded using ACF to 50-µm by 50-µm pads on the polyimide package. For the 350-µm-thick transducers, a pocket is created in the polyimide package (see **Supplementary Discussion S2**). Exposed conductive interfaces are passivated with a biocompatible epoxy.

### Demonstration of *in vivo* PNS stimulation in the New Zealand White Rabbit (NZWR)

To demonstrate VNI platform compatibility with conventional interventional radiological techniques, flexibility of the minimally invasive delivery procedure, and the efficacy of the VNI device *in vivo*, we implanted the VNI device into a NZWR and delivered electrical stimulation from the common carotid artery (CCA) to engage the nearby carotid sinus and vagus nerve. For these experiments, the surgical incision for device delivery is made in the right groin of the rabbit below the inguinal ligament, and both the common femoral artery (CFA) and common femoral vein (CFV) are accessed. The right CFA is used for navigation to the carotid sinus and delivery of the VNI device. For some experiments, the right CFV is used for navigation to the internal jugular vein (IJV) and placement of an electrophysiology catheter (4F, quadripolar), which senses the activity generated from the VNI device in the CCA. Finally, the left CFA is also accessed through an incision in the left groin and a blood-pressure-sensing microcatheter is placed for data collection.

For the 3.5-3.9-kg rabbits employed in these studies, the CCA is approximately 1.8-mm in diameter. Endovascular navigation to the CCA and IJV is achieved with a guide catheter system mediated by a microwire under fluoroscopic guidance using conventional techniques (see Methods), and vessels are checked for patency throughout the experiment (see **Supplementary Fig. S10**). Experiments that use an electrophysiology catheter delivered adjacent to the VNI deployment target typically have placements as shown in the fluoroscopy image of **Fig. 4a**. The commercial ATL P4-1 linear ultrasound probe is then placed in contact with the shaved neck of the rabbit with acoustic coupling gel. Baseline blood pressure and heart rate measurements are taken.

**Figure 4.**
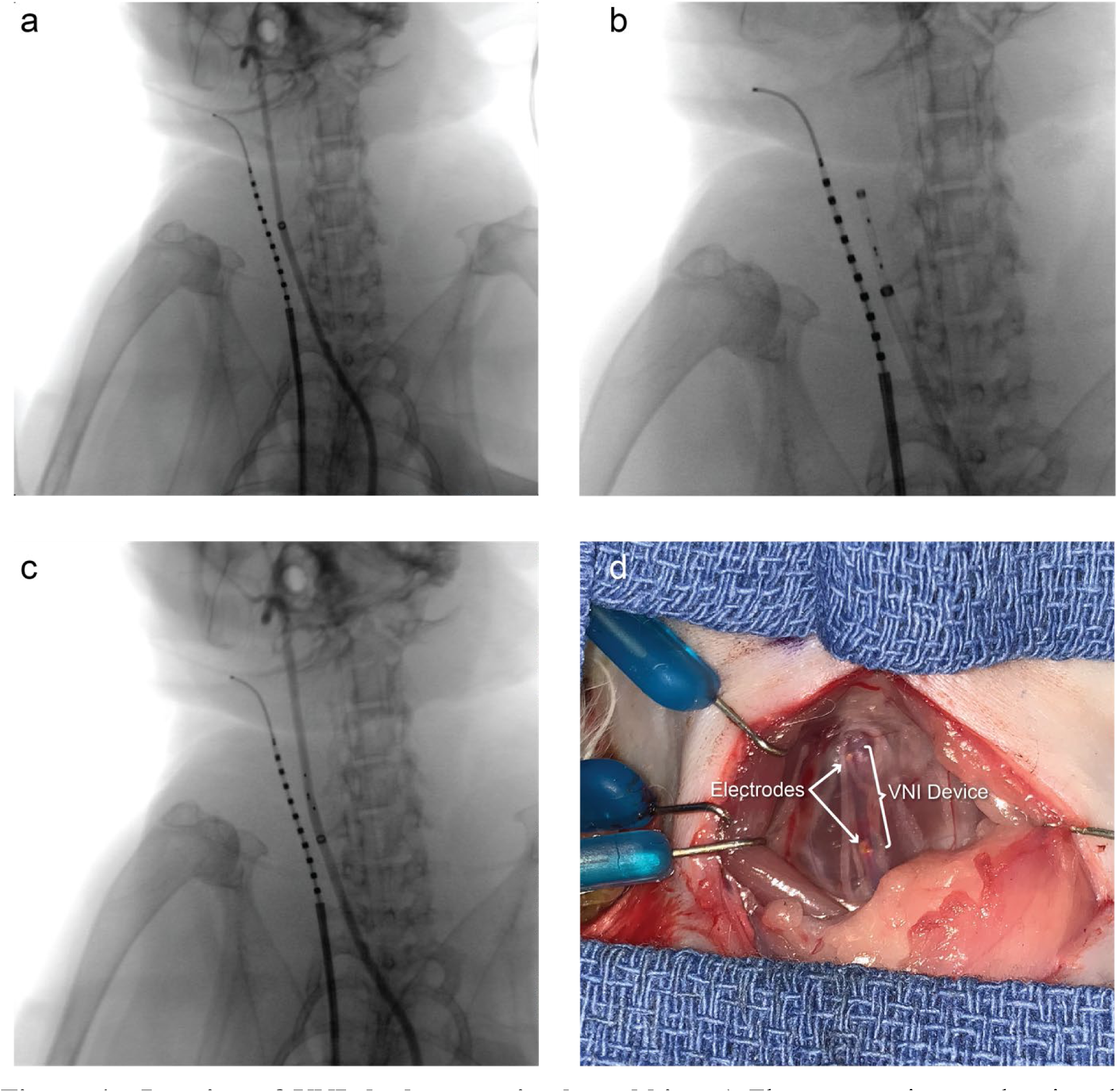
Imaging of VNI deployment in the rabbit. **a)** Fluoroscopy image showing the placement of the electrophysiological microcatheter in the IJV adjacent to catheter device deployment in the CCA. **b)** Fluoroscopic image demonstrating positioning of the VNI device at rabbit C5 cervical level immediately prior to deployment. **c)** Fluoroscopy image of the delivered VNI device with contrast, demonstrating arterial patency after insertion. **d)** Surgical dissection following the procedure in which the VNI device is shown inside the 1.8-mm rabbit CCA; the VNI device (annotated) is most easily identified by its gold electrodes (annotated).

The VNI device is rolled and inserted into the distal end (see **Supplementary Discussion S6**) of a microcatheter delivery vehicle, and is passed through the guide catheter system in the CCA to a delivery target as shown in the fluoroscopy image of **Fig. 4b**. The implant is deployed by advancing a microwire proximal to the VNI device to maintain positional stability while the delivery system is withdrawn. Retraction of the sheath permits the thin film polyimide package to unfurl and appose the vessel wall at the intended target site, in this case at the rabbit cervical C5 level, as shown in the image of **Fig. 4c**. Following insertion, the implant can be localized using the beacon response subsystem previously described.

The surgical access points, acoustic probe placement, and anatomical target for the VNI device are shown in **Fig. 5a**. Pulsed-excitation mode is used for the VNI stimulation protocol in these experiments due to the relatively low stimulation pulse repetition rate to minimize tissue heating. A VNI implant configured to deliver stimulation pulses at 300-µA-amplitude monophasic for 300-µs, with 1-ms of charge balancing occurring after stimulation current delivery. We stimulate for a total of 30-s with a repetition rate of 100-Hz. For every 10-ms period, the VNI is powered for 7-ms, providing enough time to generate the HV supply, deliver stimulation, and perform charge balancing for 1-ms, for an overall duty-cycle of 70%. An estimated peak positive pressure of 148 kPa is employed for each pulse focused with a spot size of approximately 800-µm at a depth of approximately 1.1 cm to reach the CCA. In all cases, we record the mean arterial pressure (MAP), as well as the constituent systolic and diastolic pressures, from a pressure-sensing microcatheter placed in the left femoral artery (see Methods). Some experiments also benefited from the presence of the decapolar electrophysiology catheter to record the stimulator output from the IJV.

**Figure 5.**
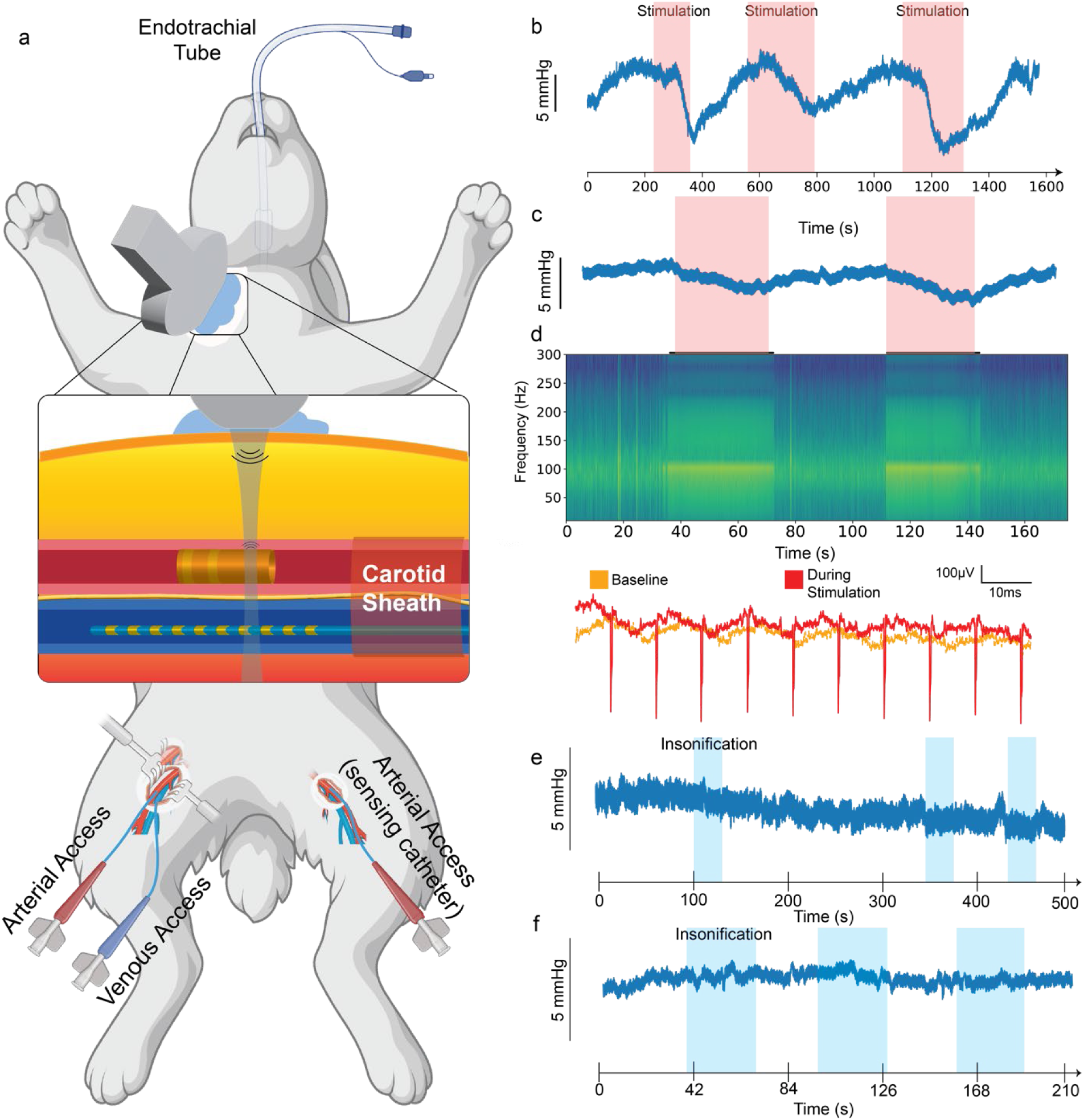
Validation of the VNI device *in vivo.* **a)** Experimental setup for the VNI experiments in the NZWR. Animal rendering is created in-part with BioRender.com. **b)** First experiment with MAP as measured from the pressure-sensing microcatheter in the left common-femoral-artery during repeated stimulation and rest periods. The VNI device is operated pulsed mode with one stimulation pulse delivered with each ultrasound pulse at a 100-Hz repetition rate. MAP data are smoothed with a one-second moving average filter. **c)** Second experiment with MAP as measured from the pressure-sensing microcatheter in the left common-femoral-artery during repeated stimulation and rest periods. MAP data are smoothed with a one-second moving average filter. The VNI device is operated pulsed mode with one stimulation pulse delivered with each ultrasound pulse at a 100-Hz repetition rate. **d)** The spectrogram shows the power spectrum of the measured electrophysiology data recorded from the IJV. Power is evident at 100 Hz and associated harmonics only during the stimulate periods. Corresponding representative time-domain traces recorded in the IJV in the absence of stimulation from the VNI device (yellow) and with stimulation activated (red) shown at higher temporal resolution. **e)** Negative control showing the lack of modulation of mean arterial pressure by the delivery of ultrasound to the CCA off focus from the location of the VNI device. **f)** Negative control from inactive VNI dummy device delivered to the CCA of the NZWR. Periods of insonification are highlighted in blue again.

**Fig. 5b** shows the MAP trace, showing a correlated drop of the MAP during stimulation epochs with a maximum reduction of 11.2% in diastolic blood pressure, averaged across all stimulation epochs, relative to the initial baseline (see Methods). The corresponding diastolic and systolic blood pressures are shown in **Supplementary Fig. S11a**, showing a preferential reduction is systolic pressure over diastolic during the stimulation epochs. We stimulate for 30-s, then allow for autonomic nervous system (ANS) recovery for 30-s by turning off the ultrasound delivery, and consequently, the VNI implant stimulator output. Three repeated stimulation and recovery phases are performed. We extract the cardiac cycles from the one baseline and three stimulation epochs, then compare the distribution of combined derivative values from the 322 cardiac cycles during stimulation to the 130 cycles during baseline together with the 263 recovery cycles between and following the three stimulation epochs, using the two-sample Kolmogorov-Smirnov test, giving a p-value of 3.79 × 10^-7^, indicative of a strong correlation between a reduction in MAP and the application of stimulation (see Methods).

**Fig. 5c** shows the MAP trace for an experiment repeated on a different animal with a different device deployed at a similar location in the CCA relative to the cervical spine but under deeper anesthesia conditions (see Methods) with the addition of direct electrophysiological recording from the IJV. A drop in the MAP in each stimulation epoch is also observed with the corresponding diastolic and system blood pressures shown in **Supplementary Fig. S11b,** showing correlated reduction in both systolic and diastolic pressure as a result of stimulation. Using the same statistical significance test employed in the analysis of **Fig. 5b** over two repeated stimulation epochs, compared against one baseline epoch we find a p value of 0.04, indicative of statistically significant correlation between a reduction in MAP and the application of stimulation (see Methods).

We show the time-correlated power spectrogram of the electrophysiological waveforms from this measurement, confirming the application of electrical stimulation from the VNI deployed in the CCA as detected from the IJV. **Fig. 5d** presents these electrophysiological recordings from the IJV in the time domain during representative epochs, showing traces in the presence and absence of stimulation from the VNI. The recorded stimulation pulses delivered at 100 Hz are concomitant with the ultrasound delivery. Each pulse has the expected characteristics of the stimulation pulse, matching the expected 300-µs pulse width for the monophasic pulse. We note that during the charge-balancing phase of the stimulation event, we do not observe a corresponding response. This response is expected to be a narrow pulse which will be very effectively filtered over the distance to the electrophysiology microcatheter in the IJV.

The two experiments in **Supplementary Fig. S11** reveal distinct blood pressure response profiles, suggestive of different underlying mechanisms. In the first experiment, stimulation produced a narrowing of pulse pressure through concurrent reduction in systolic blood pressure and rises in diastolic blood pressure, a signature pattern of baroreflex activation^20,21^. In the second experiment, pulse pressure was preserved across stimulation, baseline and recovery epochs, with the drop in MAP driven instead by peripheral vasodilation (a reduction in systemic vascular resistance), implicating preferential recruitment of efferent vagal vasomotor pathways over direct cardiac effects^22–24^.The divergence between experiments likely reflects inter-animal variability in the spatial relationship between baroreceptor afferent fibers and the main vagal trunk, as these fiber populations are known to be anatomically segregated in some species^25,26^.

Because ultrasound itself can have a neuromodulative effect, it is necessary to perform a control experiment to confirm that observed physiological responses are the result of the electrical stimulation of the device. In the first control experiment, we focus the same powering acoustic pulse train over a linear distance along the CCA of approximately 1 cm in the intended location of implant delivery but in the absence of the VNI device. The same acoustic power levels used to activate the VNI device are delivered near the CCA. Representative blood pressure recordings, shown in **Fig. 5e**, lack any acoustically mediated responses in MAP. The same statistical test applied to the stimulation experiments was applied to the control experiment after extracting the cardiac cycles from the baseline epoch against the combination of the three insonification epochs, with a p-value of 0.73, demonstrating no statistical significance. This control experiment is done prior to device insertion.

To rule out acoustic neuromodulation resulting from any acoustic effects due to the presence of the VNI device itself (i.e. not mediated by electrical stimulation from the US-powered device), we perform a second control experiment in which we implant a VNI that is mechanically identical to a fully functioning device except that the ASIC is mounted upside-down on the polyimide package, disconnecting it from the other components of the VNI. Since we lack the beacon for localization, we focus the powering acoustic pulse train on the dummy using fluoroscopy as a spatial guide for localizing the implant and the pulsatility of the CCA, discernable in b-mode imaging to determine depth. We used several different probe orientations and none resulted in a physiological response. A representative waveform is shown in **Fig. 5f**. After extracting the cardiac cycles and comparing the deltas between the one baseline epoch against the three epochs of insonification, we use the same statistical test we used in the stimulation experiments with a p-value of 0.61, indicating the changes in MAP and the insonification of the VNI device are not statistically significant.

## CONCLUSIONS

Clinical translation and widespread use of device-based therapeutics have been limited by their invasiveness, limited spatiotemporal resolution, and off-target effects. Although promising, current electroceutical neurostimulation strategies have tradeoffs between accuracy/efficacy and invasiveness. While wholly-noninvasive strategies offer an excellent safety profile, their physical distance-to-target greatly diminishes their therapeutic efficacy and stimulation accuracy ^27,28^. Focused ultrasonic stimulation (FUS) has also recently been suggested as a means of nervous system stimulation ^29^. Large acoustic intensities, the requirement for high-power beamforming arrays with specialized coupling interfaces, and complicated acoustic probe thermal management systems render this technique impractical in many cases. In this Article, we present the first fully self-contained, wireless endovascular neural stimulation system and demonstrate that miniaturization and volumetrically-efficient design allows this wirelessly-powered device to navigate tortuous structures and be successfully delivered in a small animal model with conventional surgical techniques. Our device has the potential to transform electroceutical intervention across a variety of therapies that require low-risk implantation coupled with precision therapeutic targeting.

Unlike prior endovascular devices, our implant places all electrodes and components within the vessel lumen, significantly simplifying delivery procedures and eliminating the risks and failure modes associated with other systems that require tethering that transits the vasculature. The use of a rotationally invariant system architecture reduces the complexity and requirements of endovascular delivery. Relative to traditional battery-powered implantable stimulators, wireless powering through ultrasound enables this fully-endovascular design, while also supporting a low-latency bidirectional communication link. Furthermore, by engineering a fully-self-contained device architecture, we enable delivery via other methods, such as through injection from a suitable needle (see **Supplementary Discussion S8**).

While we have engineered the initial VNI platform to fit into small-diameter vessels (< 2 mm), versions designed to operate in larger diameter vessels can benefit from reduced volumetric constraints for further optimization. In particular, the piezoelectric transducers can be used in the more common half-wave resonance mode, increasing the power available for stimulation from approximately 300-μW to 2-mW. Although this modification will increase the volume of the three piezoelectric transducers six-fold, with a resulting thickness increase from 350-µm to 900-µm, the modified VNI device could readily be accommodated in larger diameter vessels. By way of example, in a 4-mm-diameter vessel the resulting cross-sectional occlusion would be 6.4%, a modest increase compared to the occlusive cross-section of the current VNI platform.

The PMN-PT transducers used in this VNI device contain lead, and while the PDMS passivation strategy we employ in this study is suitable for acute studies, a more robust approach, such as parylene-caulked-PDMS encapsulation, could be employed to strengthen the barrier against ionic leaching ^30^. Nonetheless, there may be a preference to adopt a lead-free piezoelectric material in place of PMN-PT at the cost of available power. One such material is barium titanate, resulting in a reduction in available power of ∼90%; another option is BCZT with a reduction in power of∼60%. When scaled to operate in a half-wavelength resonance mode, however, the achievable power envelope with BCZT would be approximately the same as the existing VNI platform.

Ultrasound powering and communication offers the advantage of controllably directing energy to more than one device through acoustic beamforming. Additionally, multiple stimulation channels could be supported by a single device; ring electrodes could be partitioned into individually addressable arc-length segments, providing for more precise field shaping of the electrical stimulation. Other electrode materials such as sputtered iridium oxide (SIROF) ^31^, glassy carbon^32^, platinum black^33^, and polymeric coatings such as PEDOT:PSS^34^ could bring impedance advantages over the gold electrodes employed here.

Using an electrophysiology catheter as a positive control provides operational confirmation in the acute demonstrations presented here; however, this strategy is not viable for chronic experiments or clinical translation. Currently we employ LSK backscatter communication to transmit a single bit back to the acoustic probe for localization; however, the data payload could be expanded to include additional diagnostic information, such as the amount of charge delivered in a stimulation pulse, the stimulator voltage supply, and other similar metrics of system health. LSK backscatter can also allow a device that records neural activity or senses biomarkers in the blood.

CNS endovascular intervention is also possible with these devices and may be attractive due to the chronic stable interface formed by endovascularly deployed electrodes. The device performance of directly implanted electrodes in the brain degrades over time due to the foreign-body responses^35,36^, resulting in the need to continually increase stimulation power to maintain a consistent therapeutic effect. The fully self-contained VNI device may be particularly attractive in applications requiring minimally invasive delivery to the CNS with scale beyond existing solutions. One critical barrier for these applications is transmitting ultrasound through the skull, which presents dual challenges with both significant acoustic attenuation (∼20 dB/cm·MHz) and large interfacial reflections due to the acoustic impedance mismatch between the skull (∼20 dB/cm·Mhz) and soft tissue (∼1.8 dB/cm·MHz). Nevertheless, transcranial ultrasound has been shown to be safe, with demonstrations of focused power delivery at a focal depth of 4.5 cm with an ultrasound carrier frequency of 2 MHz ^37^, opening the door to accessing deep structures of the brain through vascular navigation.

In addition to endovascular deployment, delivery in the subarachnoid or inside cerebral ventricles is also possible, as is deployment in other catheter-accessible hollow organs such as the genitourinary or gastrointestinal tracts. This versality significantly expands the potential reach of the VNI platform, enabling access to a wide range of neuromodulation targets that were previously difficult or impossible to reach using conventional surgical or percutaneous approaches. By leveraging existing anatomical conduits, the platform may allow for minimally invasive modulation of both central and peripheral neural structures, ultimately broadening the clinical applications of neuromodulation across multiple organ systems.

## ACKNOWLEDGEMENTS

KLS acknowledges support from the Defense Advanced Research Projects Agency under Cooperative Agreement D20AC00004. ESB gratefully acknowledges support from Lisa Yang, John Doerr, Lore McGovern, the Howard Hughes Medical Institute, the Kavli Dream Team, the Ludwig Foundation, and the National Institutes of Health under R01MH117063 and 1R24MH106075.

The authors would like to thank Eric Leuthardt, Robert Desimone, Nir Grossman, Adam Marblestone, Christian Wentz, and Elazer Edelman for helpful discussions and Harbaljit Sohal, Ilke Uguz, and Jason Fabbri for fabrication assistance.

## AUTHOR CONTRIBUTIONS

JWS, GTF, ESB, and KLS conceived the project. JWS designed the implant circuit, packaging, firmware and software. GTF developed the self-expanding polymeric implant concept,. GTF, EFS, JH, JWS, and DO, developed the VNI surgical delivery and experimental protocol. JWS developed the ultrasound techniques employed. GTF, JH, DO, and JWS performed *in vivo* experiments at the Massachusetts Institute of Technology. JWS, GTF, EFS, and JS performed *in vivo* experiments at Columbia University. JH and MJ provided essential support for all the rabbit studies carried out at the Massachusetts Institute of Technology. EZ and AR provided essential support for all the rabbit studies carried out at Columbia University. JWS and GTF analyzed the data. KLS, ESB, EK, SC, and SL provided supervision. ESB and KLS provided funding support. JWS, GTF, EFS, and KLS wrote the manuscript with contributions by all authors.

## DATA AVAILABILITY

All recorded electrophysiological data relevant to the figures presented in this paper are available at https://github.com/klshepard/vni. All other relevant data are available from the corresponding authors upon reasonable request.

## CODE AVAILABILITY

All scripts used for data analysis are available at https://github.com/klshepard/vni. All other relevant codes are available from the corresponding authors upon reasonable request.

## SUPPORTING INFORMATION

Supporting information includes Discussions S1 through S8 and Figures S1 through S11.

## MATERIALS AND METHODS

### Design and Fabrication of the Application-Specific Integrated Circuit

The VNI ASIC was designed in a custom mixed-signal design flow. The generated GDSII design format was validated against design rules specified by the foundry using Calibre Design Rule Check (nmDRC, Siemens EDA) and Calibre Layout-vs-Schematic (nmLVS, Simens EDA). Further validation of the layout was performed using Calibre Resistance-Capacitance Extraction (xRC, Siemens EDA) to generate netlists that include parasitic RC components to more accurately account for the non-idealities in nanofabricated layouts. The system controller and other digital systems are written in System Verilog hardware description language (HDL) and simulated using NC Verilog (Cadence). The netlists are synthesized using Design Compiler (Synopsys), and undergo place and route using Innovus (Cadence). The digital logic is validated at the gate level and the place and route stage using the same test harnesses used for validation in the initial behavioral Verilog to ensure functionality. A top-level simulation is performed using Virtuoso ADE (Cadence) and the Spectre (Cadence) simulation engine for final checking and verification. The integrated circuit is fabricated in a 0.18-µm mixed-signal by the Taiwan Semiconductor Manufacturing Company (TSMC) in a 1.8V-5V 0.18-µm mixed-signal, radio-frequency, general-purpose (MSRFG) process.

### Design and Fabrication of the VNI

Device fabrication includes the fabrication of the flexible polymeric stent with integrated electrodes; the attachment of the electronic components to this stent including the integrated circuit, piezoelectric transducers, and energy storage capacitor; and the biocompatible encapsulation of these components. The fabrication flow is shown in **Supplementary Section S8**.

A four-inch single-side polished (SSP) silicon wafer is selected as a carrier for the polymeric stent fabrication. A layer of PI2611 is spincast on the silicon wafer at 3000 rpm. This film is soft baked at 120 °C for 3 minutes, then cured at 300 °C in a nitrogen-rich environment for 60 minutes. An 18nm/200nm titanium/gold metal interconnect layer is then patterned on the flexible polymeric stent utilizing AZ5214-E image reversal photoresist and an Angstrom Deposition electron-beam evaporation tool. After lift-off, a passivation layer of polyimide, PI2610, is then spun on at 4500 rpm and cured in a nitrogen-rich environment for 80 minutes at 300 °C for a resulting total substrate thickness of 7 µm. The interconnect layer contains metal pads corresponding to the integrated circuit pads; a 10 µm layer of P4620 photoresist is lithographically defined as a mask to etch through the passivation layer to the interconnect layer using reactive ion etching. To ensure robust electrical connection to the underlying metal layer, 1-µm-thick copper pillars are deposited via sputtering using an AJA Orion Sputtering deposition system to provide via metal from the package to the IC pad structures.

The package is then flipped over for electrode fabrication. A 10-µm-thick layer of P4620 photoresist is again photolithographically patterned to etch vias through the substrate to the interconnect layer for the electrodes. Then a secondary layer of 8nm/200nm titanium/gold is deposited to form the electrodes. The substrate is flipped back over for flip chip bonding. An 8-µm-thick HighTech TFA22020 anisotropic conducting film (ACF) with 5 µm conductive balls is applied to the substrate over the padframe. The ACF provides the adhesive underfill and conductive interface to hold the chip in place. A Finetech Fineplacer Lambda tool is used to flip-chip position the IC over the pads of the package and perform bonding by applying a force of 20N and heating to 180°C for 80s.

A pocket for the piezoelectric transducer is ablated using an IPG IX-255 excimer laser. A bulk sheet of the 350-µm-thick PMN-PT transducer (TRS Technologies, Inc., State College, PA) is cut to dimensions of 350 µm by 770 µm (oriented along the length of the vessel wall in-line with the IC) with a DISCO DAD3220 dicing saw, and is mounted to the package with a thin layer of H20E low temperature conductive epoxy. An 0201 low-profile (200-µm thick) 100-nF decoupling cap is affixed to the package using H20E low temperature conductive epoxy. A thin layer of polydimethylsiloxane (PDMS) is used to encapsulate and passivate the transducer, capacitor, and the chip.

### Rabbit Surgical Procedure, Anesthesia and Surgical Approach

All animal procedures were carried out in full compliance with applicable federal, state, local, and institutional guidelines for the use of animals in research. The procedures were approved by the Institutional Animal Care and Use Committee (IACUC) of Columbia University (CU) and of the Massachusetts Institute of Technology (MIT), under protocols AC-AABI3611 “Active stents for stimulation of the peripheral nervous system” (CU) and 0113-008-16, 1115-111-18, 1218-100-21, 1221-104-24 ‘Principles and applications of neuroscience technologies” (MIT). Male and female 3.5 – 3.9 kg New Zealand White Rabbits (Crl:KBL(NZW)) were purchased from Charles River Laboratories, MA, and maintained in AAALAC-accredited animal facilities under controlled environmental conditions until employed in the studies.

All rabbits underwent general anesthesia using mechanical ventilation with continuous hemodynamic monitoring. A 22G catheter secured in a marginal ear vein allowed intraoperative support fluid (normal saline) and drug administration. Animals were not fasted for food or water prior to anesthesia.

Anesthesia was induced by intravenous administration of propofol (2-6mg/kg), midazolam (0.1-0.5 mg/kg), and inhaled 3-5% isoflurane delivered in oxygen at a rate of 0.8 L/min. After endotracheal intubation with a 3.0 mm cuffed endotracheal tube, mechanical ventilation was initiated and the anesthesia level reduced to 0.5-2.5% isoflurane in oxygen, supplemented with a continuous infusion of intravenous ketamine (0.5-5mg/kg/hr) and fentanyl (0.005-0.05mg/kg/hr) for maintenance in some experiments.

The rabbits are placed in a supine position. Bilateral groins and neck are prepped and shaved. Lidocaine (2%, 2 mL/injection, Covetrus) is injected subcutaneously along the incision line for bilateral common femoral artery (CFA) and right common femoral vein (CFV) access. A 2-cm diagonal incision along the plane of the inguinal ligament is made, the subcutaneous fascia is retracted revealing the adductor magnus medially and the vastus medialis laterally. Dissection towards the inguinal ligament reveals the right CFV which is tied off distally and accessed proximally using a 21-gauge single wall needle. Upon the return of venous blood, a 0.018” microwire is inserted through the needle and advanced into the CFV followed by exchange of the needle for a 4 French dilator (Micropuncture ® Introducer Set). The microwire is exchanged for a 0.035’’ guidewire (Cook G01431 Bentson Cerebral Wire Guide .035" x 180cm) and the dilator is then removed and exchanged for a 5F intravascular sheath (PINNACLE® Introducer Sheath, 5F, 10cm). Attention is then turned to the right CFA, located just underneath the cannulated CFV, which is tied off distally. A loose suture is placed around the proximal CFA which is temporarily occluded with an aneurism micro-clip. Micro-scissors are used to puncture the CFA proximally, a 4F dilator is introduced and the temporary clip is removed allowing an 0.018” microwire and introducer to advance into the iliac artery and ultimately be exchanged for a 5-French or 6-French sheath (Terumo^TM^ Glidesheath Slender^TM^ 5F or 6F, 10cm) upon confirmation of arterial backflow. A 4F sheath (Terumo^TM^ Pinnacle® Introducer Sheath, 4F, 6cm) is also inserted in an identical fashion into the contralateral CFA, and a pressure sensor catheter (Millar Mikro-Tip® SPR-524, 3.5F, or Millar Mikro-Tip® SPR-320 2F) is advanced into the sheath for continuous blood pressure recording. Prior to insertion, the pressure-sensing microcatheter is calibrated using a precision sphygmomanometer using a two-point calibration from 30-mmHg to 80-mmHg. All three sheaths are secured to the skin and perfused with heparinized saline throughout the procedure.

The signal from the pressure-sensing microcatheter is relayed through a single channel AD Instruments Bridge Amp (AD Instruments, Sydney, Australia) to a PowerLab data acquisition (AD Instruments, Sydney, Australia) unit, and the data are recorded in the PowerLab’s LabChart 8.1 software.

### Electrophysiology Catheter and Wireless VNI Device Deployment

For the catheterization and electrophysiology portion of the experiment, the rabbit anesthesia is lightened and ventilation is switched from VCV to spontaneous pressure assisted ventilation (PAV) until steady state respiratory and hemodynamic condition are reached. To this end, propofol infusion is turned off, and isoflurane is reduced to 0.5-1.5%, supplemented at CU with a continuous infusion of intravenous ketamine (0.5-5mg/kg/hr) and fentanyl (0.005-0.05mg/kg/hr) for maintenance, allowing the animals to spontaneously trigger the ventilator.

Under fluoroscopic guidance (GE OEC Fluorostar 7800, MIT, or 7900, CU), a 4F diagnostic catheter (Terumo^TM^ Glidecath 4F Hydrophilic Coated Catheter, 120cm Angle) telescoped inside a 5 F or 5.6/5.3F guide catheter (Stryker^TM^ AXS Catalyst® 5 Distal Access Catheter or J&J Envoy 5F Guide Catheter, respectively) is tracked over a guidewire through the right CFA sheath into the aorta and used to select the right common carotid artery (CCA). Angiographic images of the head and neck are obtained through the venous phase and used as guiding roadmaps. Under fluoroscopic roadmap guidance, a 5 F Envoy MDP guide-catheter (Codman^TM^ ENVOY® Guiding Catheter - MPD – 5 F x 90cm x .056") is advanced over 0.035” guidewire into the rabbit’s inferior vena cava (IVC), then brachiocephalic vein, to terminate into the right IJV. The guidewire is then exchanged for a 4/3.3 F decapolar electrophysiology stimulation catheter (Access Point MAP-IT 4/3.3 F decapolar fixed curve electrophysiology catheter w/guidewire tip) which is navigated into the EJV, positioned parallel to the CCA. The system is then connected to a continuous drip of heparinized normal saline (0.9% NaCl) flowing at a rate of 2-3 ccs/minute to ensure patency throughout the procedure. The decapolar catheter is connected to an Intan RHD 32-Channel recording headstage, connected to an RHD 128-Ch Recording Controller interface and communicates with the IntanRHX software. Recorded data is processed with a second-order 60-Hz IIR notch filter to attenuate background noise introduced in the operating room.

The device delivery vehicle comprised an outer catheter with an inner diameter of 0.044” (Stryker^TM^ DAC 044 Distal Access Catheter or Revive 044 Intermediate Catheter) with the VNI device loaded at its distal end, and an inner wire or microcatheter of appropriate size with its tip abutting the proximal end of the rolled VNI. The outer catheter with the VNI loaded at its tip was first navigated to the target site, either just distal to the carotid bulb, or at cervical spine level C5. The wire (e.g. a Terumo 0.035” Glidewire) was then carefully positioned with its tip abutting the proximal end of the VNI. To deliver the VNI, the outer sheath is retracted, while maintaining the inner microcatheter or wire in place, allowing the VNI to self-expand until it tightly abuts the vessel walls. Upon device delivery and deployment, the entire delivery system is removed, and the patency of the vessel is verified.

### Acoustic protocol and acoustic negative-control experiment

Parker Aquasonic 100 acoustic coupling gel is applied to the animal’s skin, and an ATL P4-1 96 element linear array acoustic probe is placed over the VNI device delivery site. The acoustic probe is connected to a Verasonics® Vantage-256 ultrasound system, which provides acoustic beamforming, pulse-echo-response capture, image reconstruction, and device power delivery through orchestra scripts written in MATLAB. The Verasonics is put into B-Mode imaging mode, and the CCA is identified visually from observing its pulsatility. The Verasonics is then put in focused power delivery mode, using the same 7-ms 2-MHz acoustic pulses with a 100-Hz repetition rate, excitation sequence as is used to program and power the VNI device. We record blood-pressure control traces in 30-s epochs using the SPR-524 pressure sensing microcatheter connected to the AD PowerLabs data acquisition unit.

### Acoustic protocol and VNI device experiment

The Verasonics is placed in a B-Mode imaging mode and the CCA is again identified from its pulsatility. We follow along the CCA, switching between B-Mode imaging and pulse-echo response identification mode to identify the VNI device. Upon determining the device location, we deliver 7-ms 2-MHz ultrasound pulses at a 100-Hz repetition rate. Each 7-ms acoustic pulse is sufficient to power up the implanted VNI device, program the implant, allow its stimulation voltage rail to stabilize, and deliver a 300 µs monophasic current pulse of 300 µA. The acoustic probe is secured with a Noga power arm or held in place by hand.

### Data processing and analysis

Arterial pressure traces are recorded over the duration of baseline, stimulation, and recovery epochs with each epoch temporally labeled. The 30-s of the arterial pressure data immediately preceding the first stimulation period are selected as the baseline data. Stimulation epochs are defined as periods during which the VNI device is powered and delivering stimulation pulses. Recovery epochs are defined as time periods between stimulation epochs along with the 30-s following the final stimulation epoch. For each epoch, cardiac cycles are isolated by finding the systolic peaks, the diastolic troughs, and calculating the mean time and mean arterial pressure (MAP) from these extracted values. The cycle-to-cycle MAP delta is calculated and used for statistical analysis to determine if the effect of the stimulator on the MAP is statistically significant. The baseline MAP delta data combined with the recovery period MAP delta data are taken together as periods during which the stimulator is not active. These data are compared against the MAP-delta data extracted from the stimulation epochs using the nonparametric two-sample Kolmogorov-Smirnov test to determine the statistical significance of the application of the VNI stimulation. Stimulation-related reductions in MAP are normalized to the percent of the baseline, and reported as a percent reduction. For MAP traces that experienced sensor DC-drift a one-second moving average filter is applied to the entire experimental trace, and the data are linearly normalized to remove the drift across the duration of the time-series measurement data while preserving the hemodynamic changes on a per-epoch basis. The trace is then normalized to the baseline systolic and diastolic pressure range measured using a pressure cuff in the operating room during the experiment. Changes occurring within the time frame between initiation of stimulation and end of recording (typically t=30s from stimulation initiation) is attributed to stimulation effect as previously described ^41–43^.

For the electrophysiology catheter recordings, electrophysiology data are recorded for baseline, stimulation, and recovery epochs and subsequently processed with a second-order IIR 60-Hz notch filter using the Python signal processing toolbox followed by a fourth-order IIR 100-Hz bandpass filter to isolate the frequency band of interest. During stimulation, the onset time is noted for both of the electrophysiology measurement and pressure acquisition, and the MAP recording presented is temporally aligned to the electrophysiology recording based on the times noted for both measurements.

