## Supplementary Information for "A Fully Endovascular Neural Interface"

**Supplementary Discussion S1. Implanted Transducer Design**

Design of the acoustic power transfer begins with the selection of the link frequency, which is a trade-off between length-scale resolution and acoustic power attenuation in soft tissue. Acoustic wavelength is determined by the acoustic phase velocity equation:

$$\lambda = \frac{c_{tissue}}{f_{us}} \quad (1)$$

where $c_{tissue}$ is approximated as 1540 m/s for fluid and soft tissue and $f_{us}$ is the ultrasound operating frequency. Acoustic attenuation properties of soft tissue increase linearly with ultrasonic frequency. For example, the attenuation coefficient $\alpha$ for adipose tissue is, 0.29 $\frac{dB}{cm \cdot MHz}$ [1]. With a target of at least 5-cm of operational depth, choosing an ultrasound frequency in the 1-2 MHz range offers a favorable trade-off, providing mm-length scale wavelengths while maintaining low attenuation characteristics in soft tissue for deep tissue implants, and reasonable focal spot-sizes for implants shallower than the focal depth limit. After establishing a suitable frequency range, we turn our attention to the design of the implanted receiver. Most recently proposed acoustically powered systems utilize $mm^3$ PZT based transducers to receive power, however this is impractical for the VNI design. A 1-mm thick piezoelectric transducer would extend the VNI device more than halfway into the target vessel, occluding the cross-section of the target vessel diameters by 33% from the piezoelectric material alone. This strategy becomes particularly untenable once the piezo mounting and passivation packaging are added to the occluding cross-section in the vessel. Furthermore, loading the VNI device in suitably sized delivery microcatheters would be intractable. To achieve low-volume transducer dimensions while maintaining high charge-availability, we utilize the piezoelectric PMN-PT material, which has a favorable electromechanical coupling coefficient ($k_{33} \approx 0.93$), and exceptional piezoelectric charge coefficient in the longitudinal dimension ($d_{33} \approx -1250\, pC/N$) [cite]. While designing the dimensions of the transducer, it is advantageous to design the transducer to operate with a predominantly real impedance, occurring at open circuit and short circuit resonances, which can be determined from the following [2, 3]:

$$f_{oc} = \frac{v_p}{2t} \quad (2) \qquad\qquad f_{sc} = \frac{f_{oc}}{\sqrt{1+\frac{8}{\pi^2}\frac{k_{33}}{1-k_{33}^2}}} \quad (3)$$

Here, $v_p$ is the speed of sound through the transducer material, $t$ is the thickness of the transducer, and $k_{33}$ is the electromechanical coupling factor discussed above. Based on this, a ~2 MHz short-circuit resonance can be achieved with 350-µm thick PMN-PT. The lateral dimension was chosen to be the same as the integrated circuit width (350 µm), and the length of the transducer was set to the spot size of ultrasound in soft tissue at 2 MHz (770 µm). We anticipate that the measured resonant frequencies will be lower than the calculated frequency, due to higher vibrational modes in the non-cubic material (i.e., l >> w,t). As a result, PMN-PT material (TRS Technologies, State College, PA) was purchased, as seen in **Fig. S1a**, and diced to the desired dimensions using a DAD3230 dicing saw, shown in **Fig. S1b,c**. After packaging (see **Supplementary Discussion S2**), the impedance was measured using an Analog Discovery Impedance Analyzer system (Digilent, Pullman WA), with the resulting characteristic shown in **Fig. S1d**. The short-circuit resonance was determined to be approximately 1.88 MHz, close to the selected 2-MHz operating frequency.

**Supplementary Discussion S2. Transducer Packaging Approach**

Unlike conventional piezoelectric bonding approaches, which require fragile wirebonds, substrate temperatures that can exceed 100°C during processing, and necessarily large volume encapsulation schemes, as shown in **Fig. S2a**, the VNI minimizes the overhead on piezoelectric interconnect through microfabrication of the flexible polyimide packaging, as shown in **Fig. S2b**. The piezoelectric transducers are affixed to the VNI polymeric substrate through Epotek H20E low-temperature conductive silver epoxy. A tongue structure is fabricated by laser micromachining a portion of horseshoe pattern out of the substrate, using an excimer laser with a beam-spot size of 10-μm, as shown in **Fig. S2c**. When the tongue structure is lifted, a pocket in the packaging is formed, into which the piezo can be inserted as shown in **Fig. S2c** and **Fig. S2d**. As PMN-PT has a lower curie temperature ($T_c$ of ~100°C) than other conventionally used piezoelectric materials, we establish a thermal envelope of 80°C to provide some margin to avoid depolarization of the piezoelectric material. A bulk piece of PMN-PT undergoes the same thermal process exposure and is measured before and after the epoxy curing process to ensure the material does not lose its piezoelectricity. An image of the packaged encapsulated PMN-PT transducer is shown in **Fig S2e**. The $d_{33}$ of the piece of the bulk PMN-PT process monitor material was measured using a Piezotest $d_{33}$ meter (Piezotest Pte. Ltd., Singapore), as shown in **Fig. S2f**, with an average value of $d_{33}$=-1180 pC/N before thermal processing and a $d_{33}$=-1224 pC/N after thermal processing.

**Supplementary Discussion S3. Acoustic Angular Insensitivity**

The structural VNI device is a flexible polymer that self-expands into the vascular structure in which it is delivered. Based on the spatial dimensions of the piezoelectric transducers discussed in S1, the rotational orientation of the piezoelectric positioning is impossible to deconvolve from the fluoroscopic imaging alone. Piezoelectric transducers optimally receive power when positioned normal to the incident acoustic energy, with energy transmission reducing as a function of the cosine of the angular offset for the designed vibrational mode, thus orienting the transducer for power transfer is critical for high-power applications like stimulation. Furthermore, the self-expansion of the VNI is a random mechanical process, so even if perfect alignment was achievable prior to delivery from the microcatheter the transducer may still ultimately deploy to an off-angle position as the polymeric stent scaffolding unfurls to the vessel extents.

In a practical setting, the physician has the freedom to reposition the acoustic probe, as most raylines from the skin to an *in vivo* endpoint occur over curvilinear topologies. This applies to our demonstration of the VNI technology, as we are positioning the probe on the neck to a target inside the CCA. Combining this most applications can achieve close to optimal power delivery. The piezoelectric transducers are spaced along the length of the VNI 800-μm apart to ensure vessel occlusion is never greater than 5%. Each transducer has its own electrical interface to prevent off-angle transducers from electrically loading the transducer harvesting energy. The layout of the components on the flexible polyimide scaffolding reflecting these design constraints is shown in **Fig. S3a**.

To accommodate these scenarios, we place a total of three piezoelectric transducers on the VNI device, laid out to be spatially offset around the vessel by 120 degrees. The transducers are designed to receive power from either side, providing a maximum of 60 degrees of offset delta, or a maximum of -3dB of power loss due to angular offset. A three-dimensional mockup is shown in **Fig. S3b**. We characterize the rotational invariance of the VNI with a stationary unfocused acoustic probe, rotating the implant, through 2cm of mineral oil and the power delivery set to 14% of the FDA limit. The test setup is shown in **Fig. S3c**. The resulting rectifier voltage curve is shown in **Fig. S3d**, demonstrating that for shallow implants the VNI can be powered to a viable level without repositioning the acoustic probe, provided the implant is in the field of view of the probe.

**Supplementary Discussion S4. Circuit Architecture**

Power delivered to any of the three transducers, as discussed in **Section S3**, is rectified using a corresponding active diode rectifier. Each rectifier comparator is used to recover a clock from the acoustic pulses, which is divided by two providing a 1-MHz system clock. An activity detector monitors the three active rectifier and recovered clocks, each with a corresponding five-bit binary counters, summing clock pulse counts generated from each of the three rectifiers. The first active rectifier/piezo pair to reach 16 counts is selected as the system clock source. This prevents extraneous ultrasound reflections from generating clock glitches, although acoustic energy harvested by the off-angle transducers can still be used to augment overall system power.

An LDO regulator generates a 1.2-V logic supply. As the LDO voltage stabilizes, a power on reset circuit resets the logic and the controller transmits the beacon pulse LSK modulation. Shorting the piezoelectric transducers prevents acoustic energy harvest, and also shuts down the clocking mechanism, so a watchdog timer disables the LSK FET after a 10-µs pulsewidth. Backscattered ultrasound pulses modulate the load impedance of the piezoelectric transmitter ($|\Gamma| \propto \frac{R_{eq}}{R_{eq}+R_S}$). The system logic then enters a wait-for-programming state. Downlink data is encoded through pulse-width-modulation on the incident ultrasound pulse, with pulses greater than 16 cycles decoded as a logic '1' and pulses less than 16 cycles decoded as a logic '0'. There is a 6-µs pause in between incident acoustic pulse trains, denoting the transition between bits, which is detected with another watchdog timer. Twenty-seven control bits sent to the VNI ASIC contain information about device mode and control settings including current amplitudes, timing, and repetition rate. The programming sequence requires less than 1 ms, ensuring that low frequency probe drift does not affect transmission. After programming, a five-stage current-starved ring VCO operating between 800 kHz and 40 MHz clocks a four-stage push-pull voltage multiplier to generate a 5V stimulation supply. Charge storage is augmented by an off-chip 100nF capacitor. The VCO frequency is dynamically modulated by the stimulation supply load and the system voltage. As the stimulation supply droops, the VCO frequency increases in an effort to compensate for the stimulator load condition, however because the VCO runs at a faster rate than the powering pulses from the external probe the charge pump could pull charge from the rectified voltage and brown-out the system. A supply monitor will throttle the clock driving the charge pump to the system clock rate if it detects the system voltage has been pulled below 1.3V.

The current stimulator consists of eight slices of an H bridge driver and three current bias tuning bits enabling an output current range from 100 µA to 800 µA. The stimulator is clocked at 1 MHz and each state timing can be programmatically assigned with 5 bit tuning resolution, as well as the stimulator repetition rate. The stimulator can be configured to deliver monophasic or biphasic current pulses. An annotated chip micrograph is shown in **Fig. S4**.

**Supplementary Discussion S5. Stimulation Electrode Characterization**

The Ti/Au microfabricated ring electrodes are designed to span the circumference of the vessel facing towards the vascular wall to minimize shunt current pathways within the fluidic blood itself. The electrodes are fabricated out of 20nm Ti/200nm Au and are 1.5mm × 6.28mm at a pitch of 3.7mm. The electrodes are laid out on the base polyimide layer, and tapered vias are etched into the backside of the 6-µm thick polyimide using a low-pressure reactive ion etch. The impedance spectrum of the electrode was measured using a CHI660D electrochemical workstation (**Fig. S5a,b**) in 1× phosphate-buffered saline (PBS) (Gibco), with corresponding magnitude and phase displayed in **Fig. S5c** and **Fig. S5d** respectively.

**Supplementary Discussion S6. VNI Loading Technique**

We have developed a rolling technique to load the flexible polymeric VNI device into the end of a microcatheter delivery vehicle as small as 4.1 Fr. with an ID of 1.12mm). A 0.5-mm diameter mandrel is machined to create flat inset approximately 0.25-mm to 0.3-mm in depth along a portion of the length of the mandrel to accommodate the volumetrically significant structural components of the VNI. The mandrel is placed on top of the VNI device, initially positioned over or adjacent to the chip, first piezo, and power storage capacitor, as shown in **Fig. S6a**. and is carefully affixed using a water-soluble non-toxic adhesive. After the adhesive dries, the stent is rolled about its lengthwise-axis around the mandrel and inserted into the end of the delivery vehicle. Polyethylene glycol (PEG) can be used as a lubricant to ensure minimal friction between the delivery vehicle and the VNI polymeric structural scaffolding. The VNI device is inserted in the end of the delivery microcatheter, as shown in **Fig. S6b.** The delivery microcatheter is continuously flushed with fresh saline, displacing the PEG and causing the adhesive to dissolve which frees the mandrel from the VNI device. After no more than 10 minutes of flushing the adhesive fully dissociates from and the mandrel is pulled from the end of the delivery vehicle, while the VNI device remains securely positioned at the end of the microcatheter. The microcatheter is typically flushed for another 10 minutes with fresh saline to ensure the adhesive is fully flushed from the delivery vehicle.

**Supplementary Discussion S7. Thermal Modeling**

The Food and Drug Administration (FDA) has set the intensity limit for spatial-peak temporal-average intensity at 7.2 $mW/mm^2$ for diagnostic US applications. We seek to operate the VNI device at below 25% of this limit. The Verasonics Vantage tool has a software constraint that can be set by the user to ensure that the maximum. To ensure that the acoustic probe is not heating to an extreme, we measure the temperature of the unloaded acoustic probe, as shown in **Fig. S7**. In this measurement, we use an FLIR camera to monitor the unloaded P4-1 acoustic probe. In this configuration, the probe should have the worst-case measured temperature at the probe surface, as nearly all of the mechanical energy is reflected off of the probe interface back into the transducer, and the only means of heat transfer is convection to the surrounding environment. The probe shows a temperature elevated above the room ambient temperature, but below normal biological temperatures.

**Supplementary Discussion S8. Injectable Delivery**

Syringe injectable delivery provides a promising minimally invasive means of delivering bioelectronic systems *in vivo* opening a new frontier to how devices can be integrated into biological structures [4]. This approach not only minimizes the risks associated with traditional surgical implants but also enables a more natural and adaptive interface with biological systems. While the devices presented in this work were delivered through conventional catheterization procedures, a variant of the VNI device optimized for rapid syringe injectable delivery could be constructed to allow for rapid delivery in an endovascular structure or interstitial cavity. As a demonstration of this capability, we use the same preparation and loading procedure described in **Supplementary Discussion S6** to load the VNI device into the end of a 12-gauge syringe, as shown in **Fig. S8**. The electrodes are clearly visible, with structural elements inside the syringe needle, identical to the methodology used for the microcatheter loading.

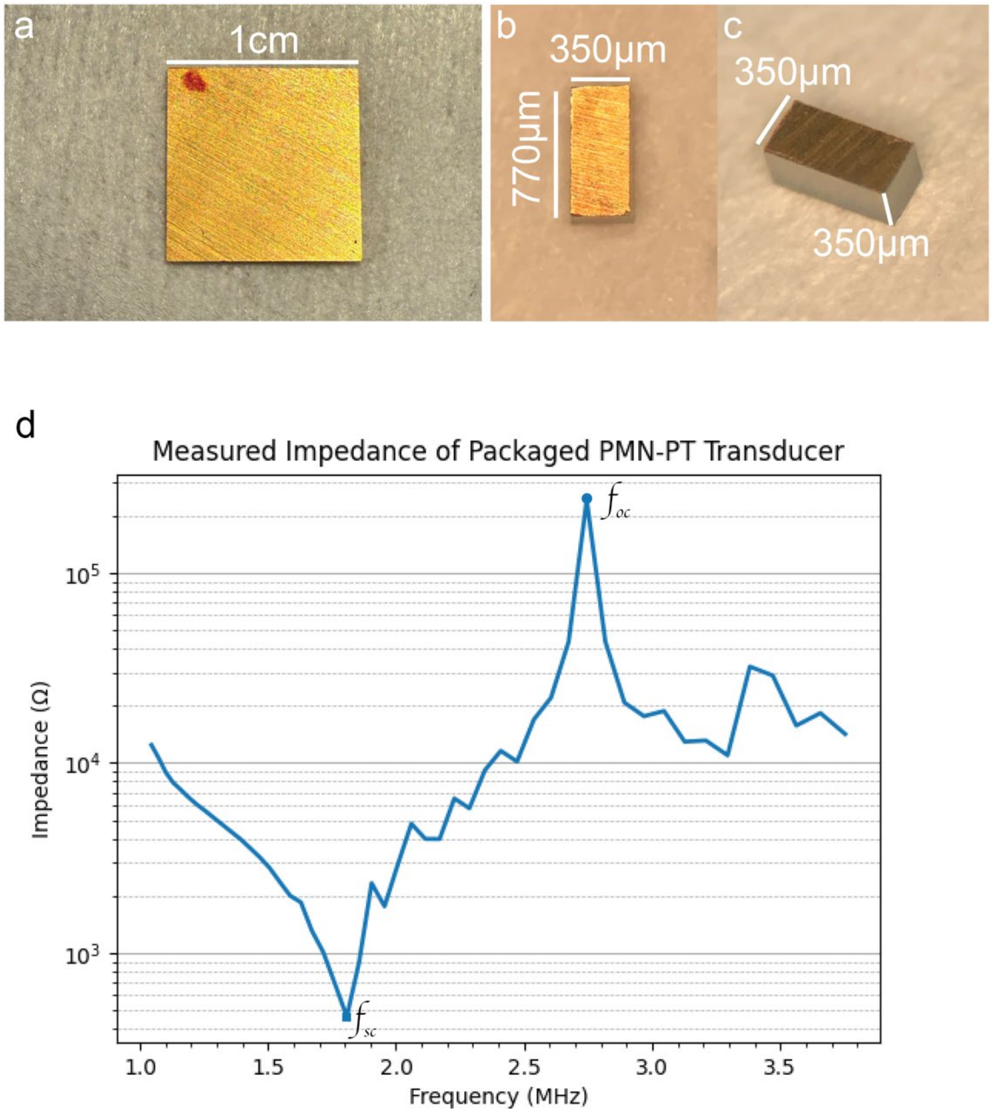


**Figure S1**. **Piezoelectric transducer design and characterization. a)** PMN-PT material as received from TRS Technologies. **b)** Top-down view of the transducer after dicing. **c)** Perspective image of the piezoelectric material after dicing. **d)** Measured impedance response of the packaged PMN-PT transducer, with short-circuit ($f_{sc}$) and open open-circuit ($f_{oc}$) resonance frequencies labeled.

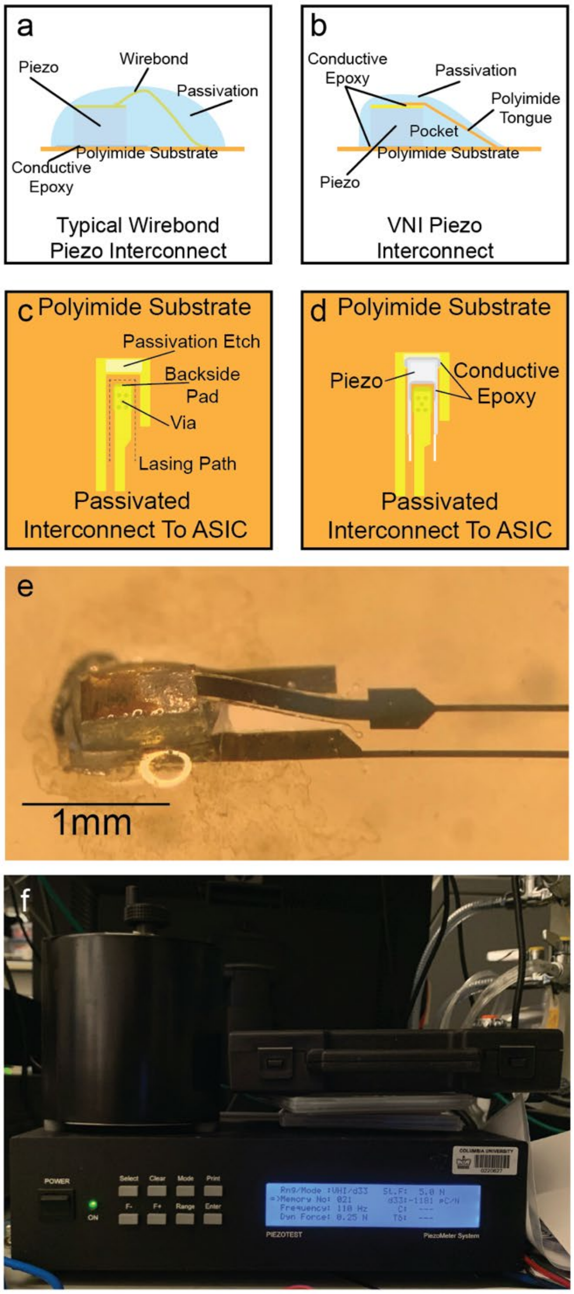


**Figure S2. VNI piezoelectric transducer packaging.** a) Schematic of conventional wirebonding approach for connecting piezoelectrics to flexible packaging. **b)** Schematic of VNI low temperature pocket approach for packaging piezoelectrics. **c)** Top-down view of package before laser ablation and mounting the transducer with low temperature conductive silver epoxy. **d)** Top-down view of mounted piezoelectric transducer before encapsulation illustrating technique. **e)** Photo of packaged and encapsulated PMN-PT transducer. **f)** Piezotest piezometer system for measuring PMN-PT $d_{33}$

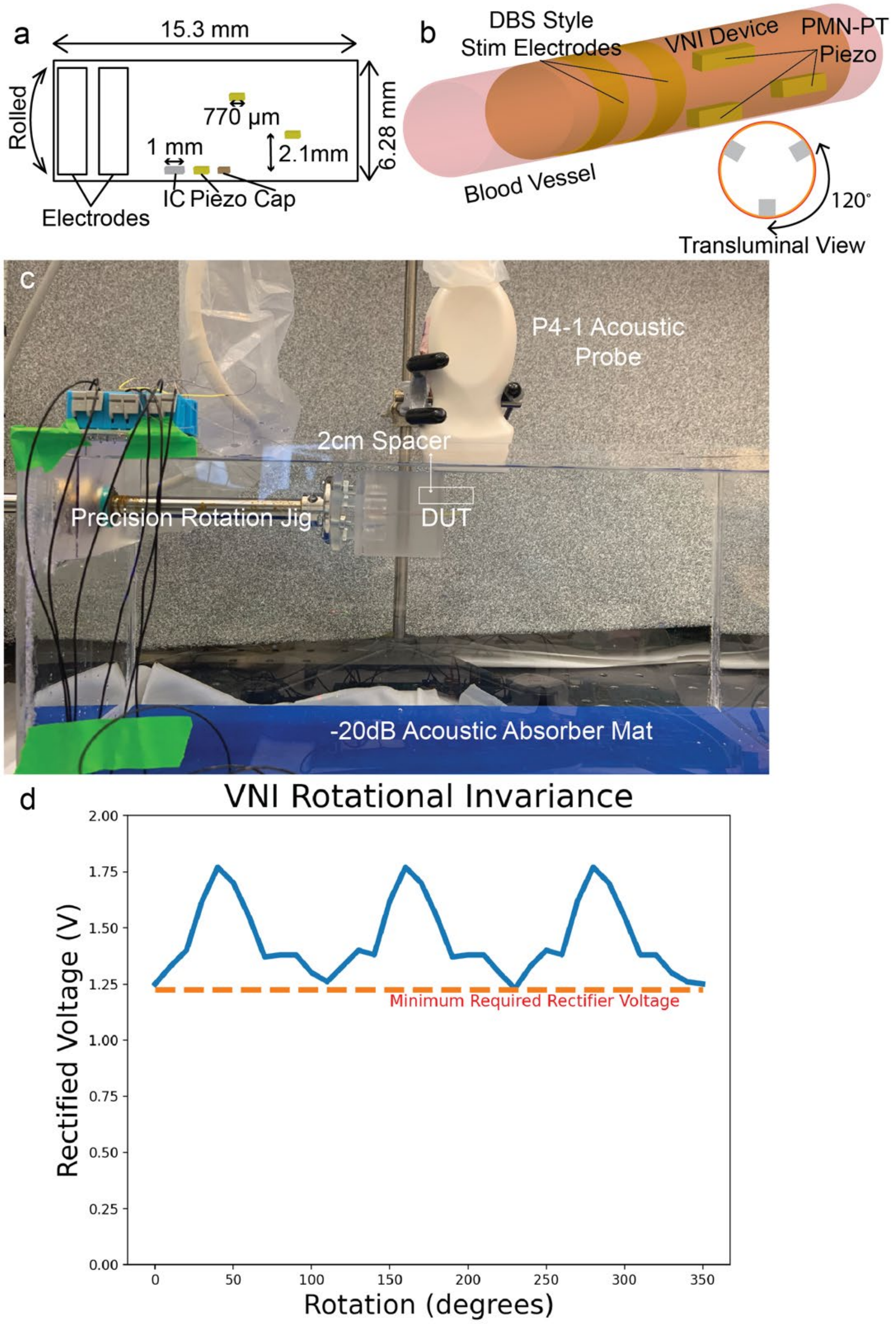


**Figure S3**. **Components related to the rotational invariance of the VNI device and the validation of operation at arbitrary insertion angle. a)** Position of the three PMN-PT piezoelectric transducers on the flat/unrolled VNI device. **b)** Position of the three PMN-PT piezoelectric transducers when rolled as in vessel deployment. **c)** Rotational test setup including spacer, position of DUT, and rotational jig. **d)** Rectified output voltage from rotational jig measurement as a function of angular offset, showing that regardless of VNI rotational orientation relative to an external acoustic probe the implant can be wirelessly powered for implantation depths of 2-cm.

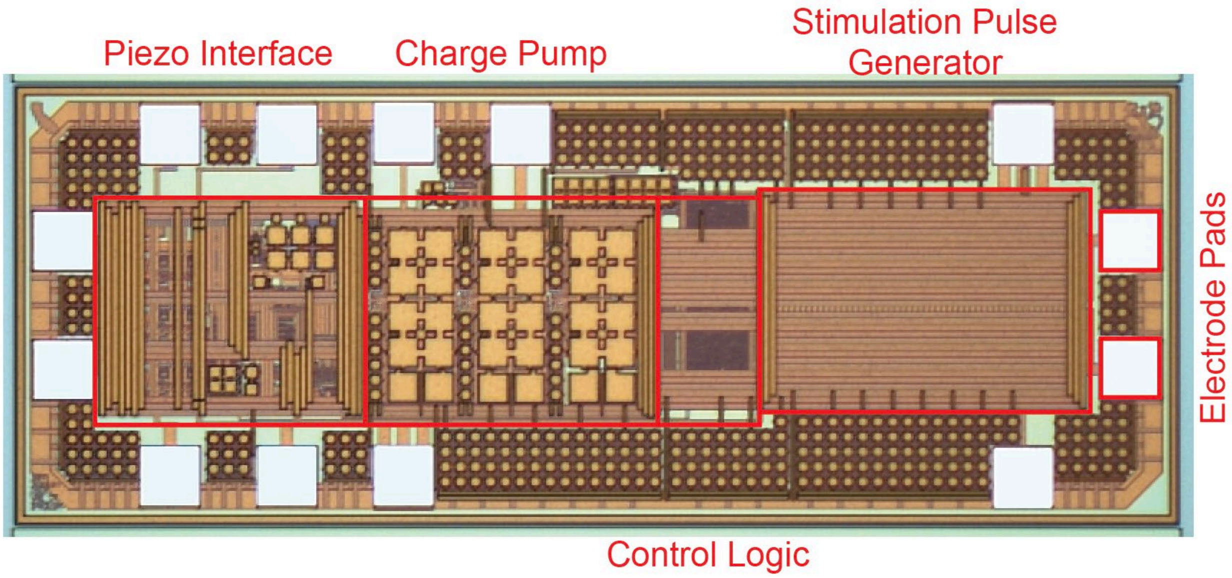


**Figure S4. Annotated die micrograph of the VNI ASIC**. The physical design and resulting implementation of each subsystem from **Fig. 1b** are highlighted in the overall VNI ASIC micrograph, and shown relative to each other. This includes the piezoelectric interface, which is responsible for rectifying incident acoustic energy transduced through the PMN-PT transducers, clock generation, data recovery, and biasing. The charge pump circuit used to generate the stimulation supply voltage. The control logic, which manages the stimulator output based on the received data sequence, and the stimulator pulse generator, which drives current into tissue in a controlled manner.

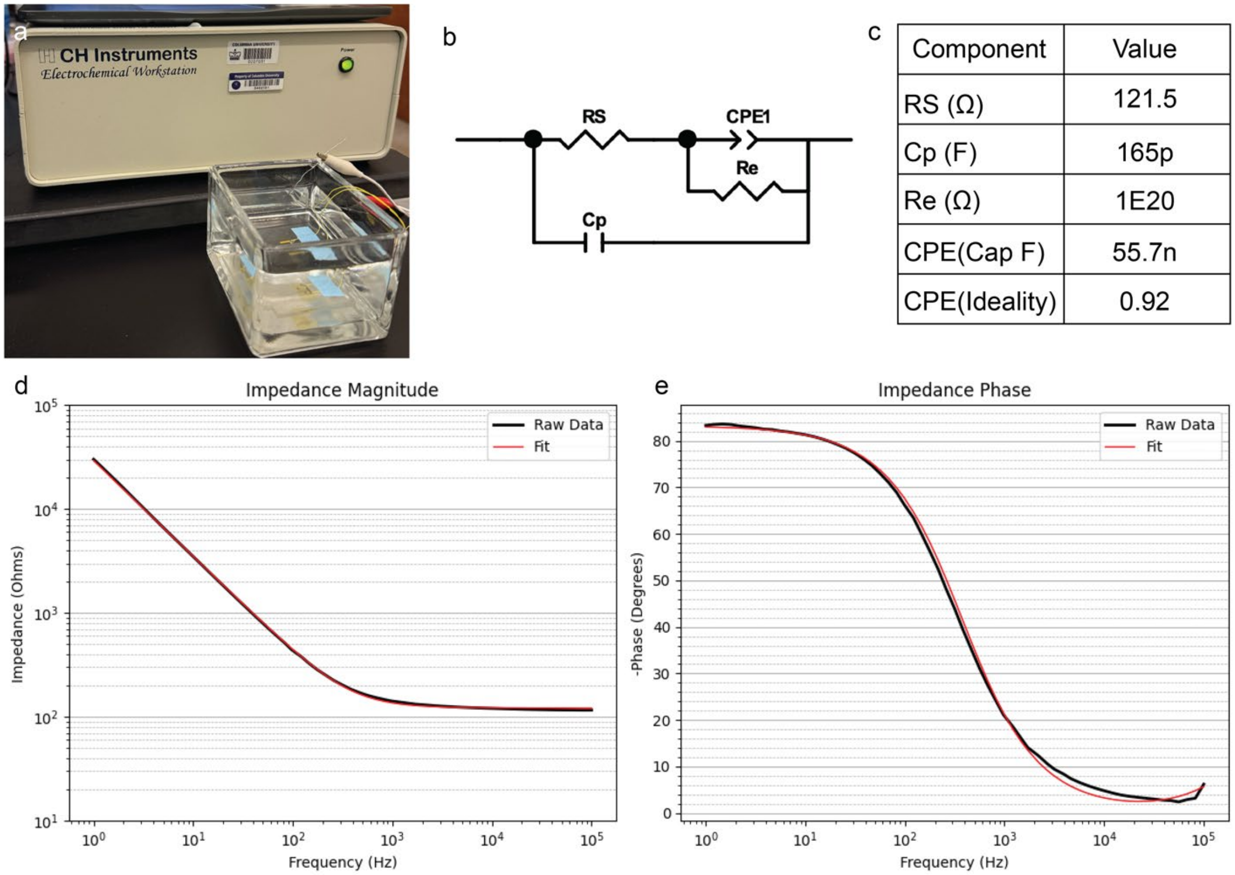


**Figure S5. Test setup and results of gold electrode electrochemical characterization**. **a)** Image of test setup, including the CH Instruments 660D electrochemical workstation connected to a glass dish with 1× PBS and VNI electrodes with passivated wiring affixed to a glass carrier. **b)** Circuit model used for fitting the equivalent magnitude and phase elements from the recorded EIS response. **c)** Fitted values for each of the components in (b). **d)** Magnitude measurements of the EIS response. .**e)** Phase response of the VNI gold electrodes measured in 1× PBS.

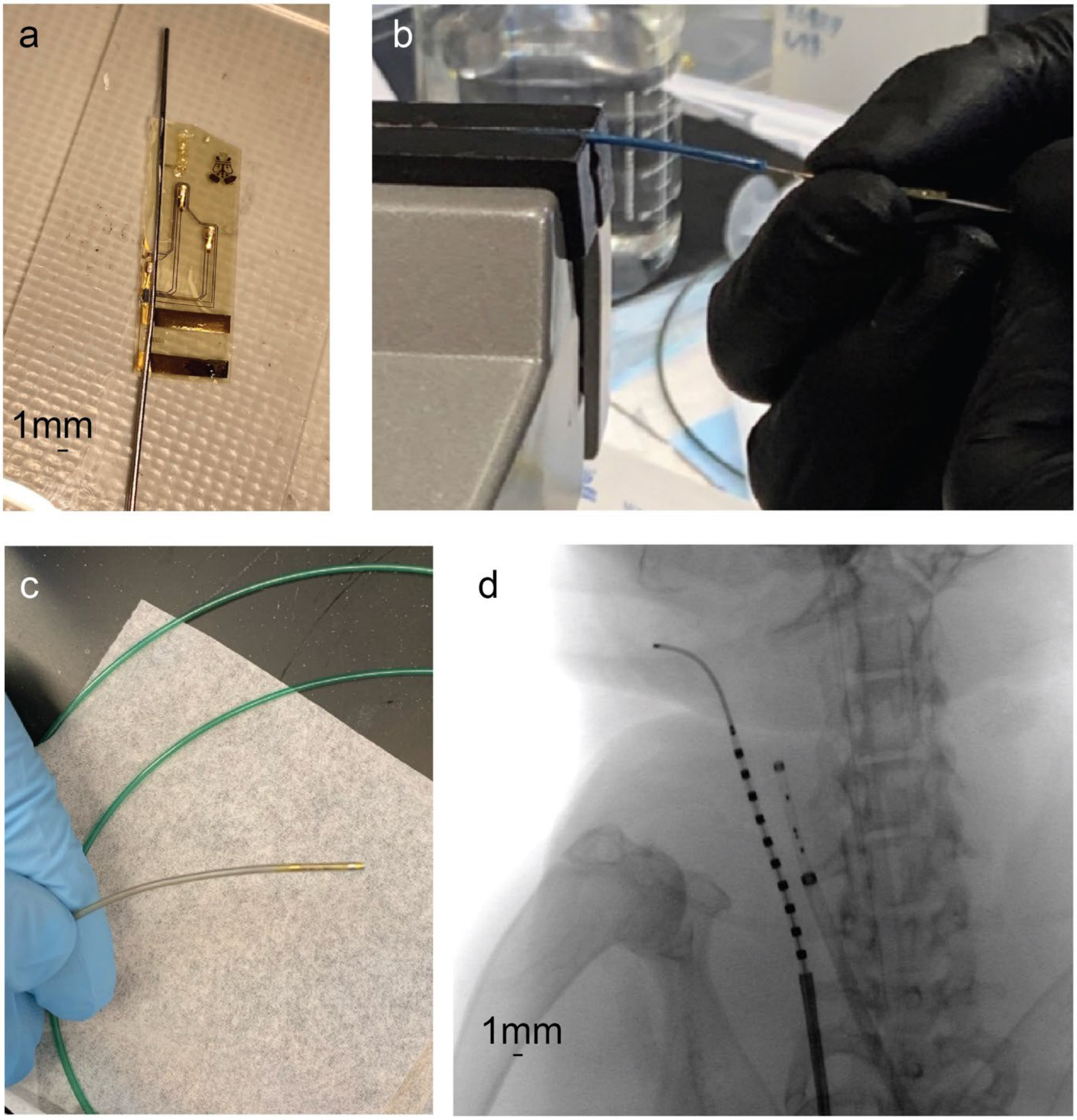


**Figure S6. Preparation of the VNI device for loading, the loading procedure, and demonstration of the loaded VNI device delivery**. **a)** Device adhered onto a mandrel. **b)** Device being loaded into microcatheter; the VNI device has been wrapped around the mandrel and is being gently positioned inside the delivery vehicle. **c)** VNI device in the end of a clear-tipped microcatheter. **d)** Fluoroscopy image of VNI device in microcatheter in the CCA of the rabbit.

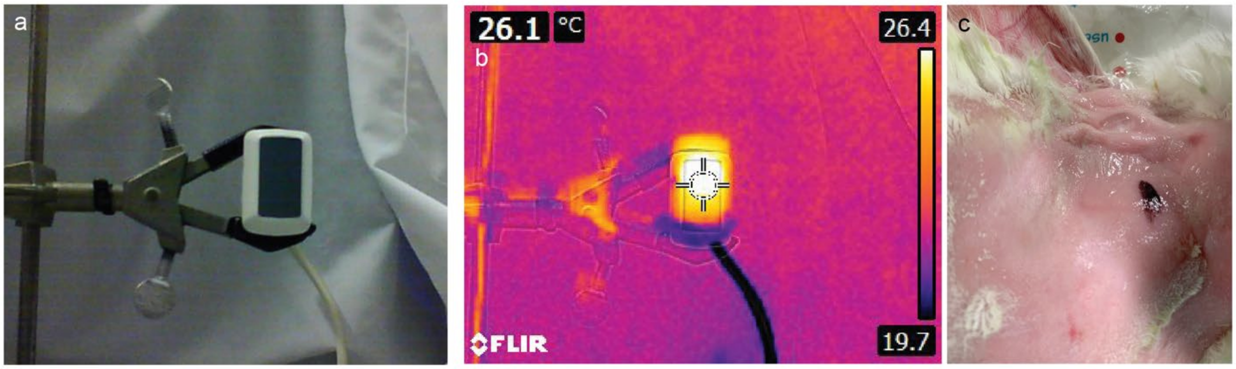


**Figure S7. Measured acoustically-induced thermal heating. a)** An image of the configuration of the P4-1 acoustic probe for this measurement. **b)** A thermal IR heatmap image taken with an FLIR camera showing relevant temperatures while the acoustic probe is energized. **c)** Image of rabbit skin after experiment termination showing the placement of the acoustic probe and no thermal damage.

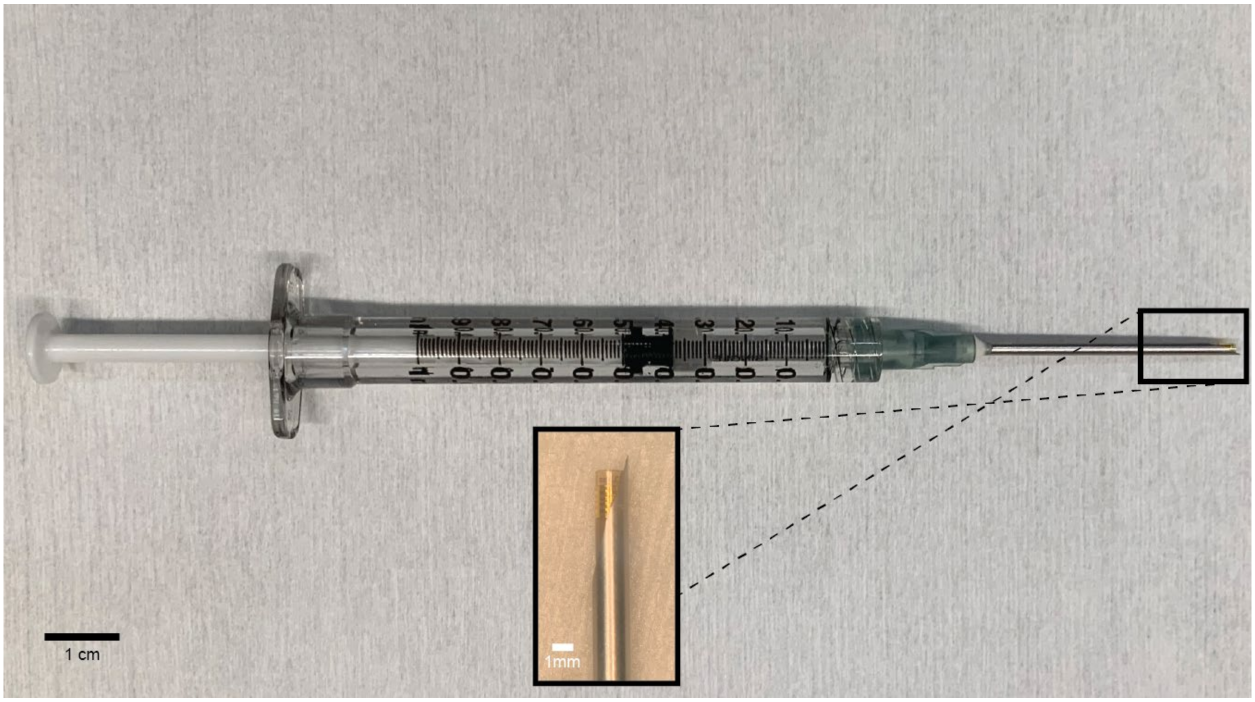


**Figure S8**. **Photo of the VNI device loaded into the end of a 12G syringe filled with phosphate buffered saline (PBS).** Close-up view shows the outer electrode of the VNI device loaded in the syringe.

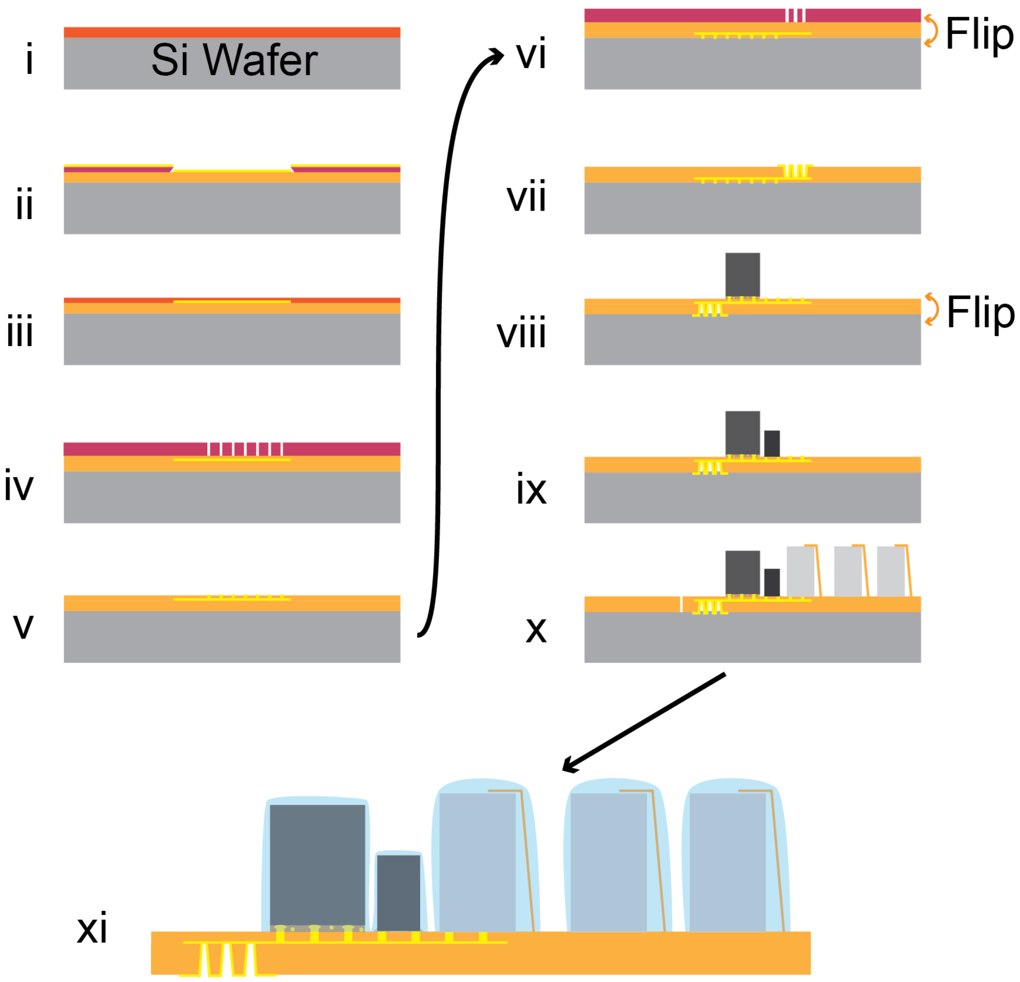


**Figure S9**. **Process fabrication flow for creating the VNI device**. **i)** Fabrication starts with selection of a four-inch single-side polished (SSP) silicon wafer used as a carrier, upon which a layer of PI2611 is spincast. **ii**) After polyimide curing, photolithography is used to define the interconnect layer, and 18nm/200nm titanium/gold metal is patterned using image reversal photolithography. **iii**) After lift-off, a passivation layer of polyimide, PI2610, is then spun. **iv)** After curing, the passivation is lithographically patterned for selective connectivity to the underlying metal **v)** To ensure robust electrical connection to the underlying metal layer, 1-µm-thick copper pillars are deposited via sputtering using an AJA Orion Sputtering deposition system to provide via metal from the package to the IC pad structures. **vi)** The package is flipped over and a via pattern to the interconnect layer is again lithographically defined. **vii)** Electrodes are deposited as 8nm/200nm titanium/gold. **viii)** The package is flipped back over and the 290-µm thick VNI ASIC is flip-chip bonded to the package with an anisotropic conducting film. **ix)** The 100-nF energy storage capacitor is bonded to the package with conductive epoxy **x)** Pockets for the piezoelectric transducer are ablated into the packaging, and PMN-PT piezoelectric transducers are affixed to the package via low-temperature epoxy. **xi)** A thin layer of polydimethylsiloxane (PDMS) is used to encapsulate and passivate the transducer, capacitor, and the chip.

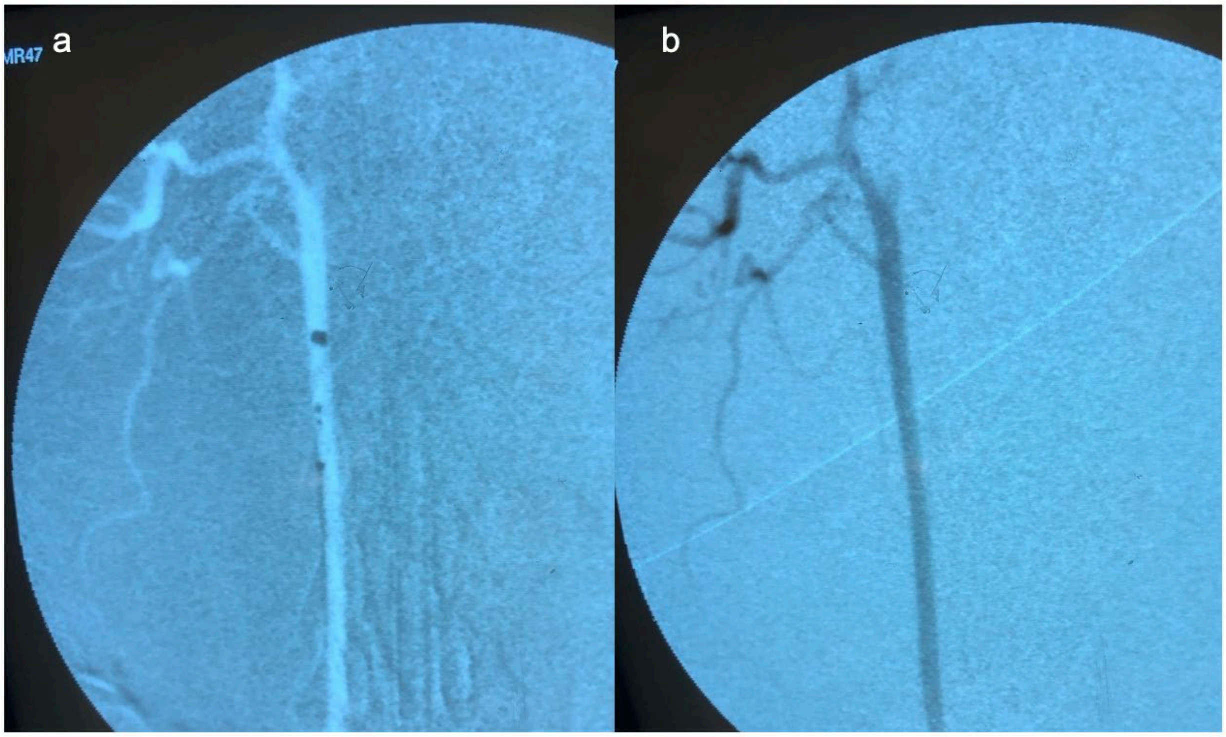


**Figure S10. Vessel patency over three hours in an acute experiment in a NZWR.** a) Fluoroscopic image showing the device in the rabbit common carotid artery during delivery. b) Fluoroscopic image following contrast agent injection showing full vessel patency retention three hours after device delivery.

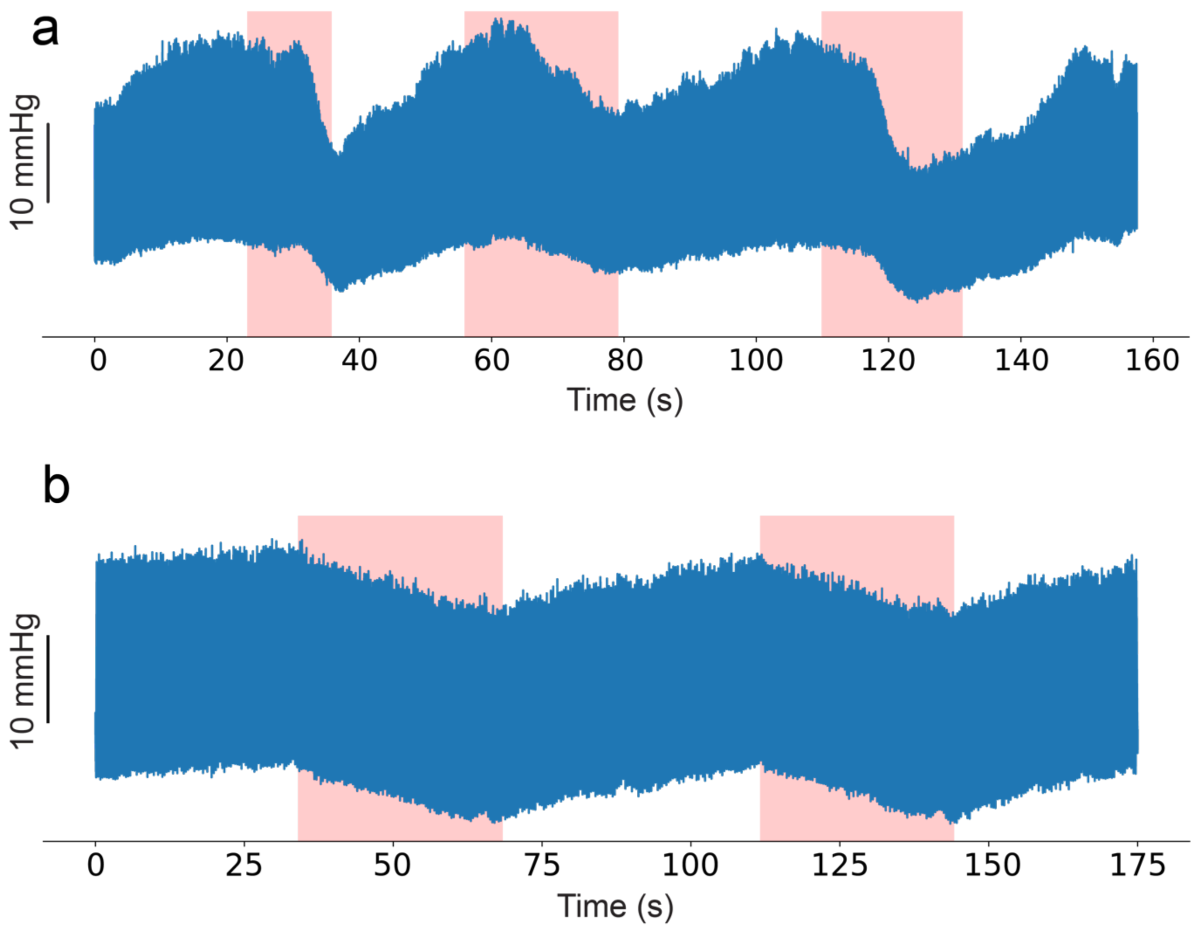


**Figure S11**. **Raw data traces of systolic and diastolic blood pressure from the two *in vivo* rabbit experiments in which stimulation epochs are noted.** a) Raw data trace from the first experiment, showing narrowing of pulse pressure through concurrent reduction in systolic blood pressure and rises in diastolic blood pressure, a signature pattern of baroreflex activation. b) Raw data trace from the second experiment, showing pulse pressure was preserved across stimulation, baseline, and recovery epochs with the drop in MAP driven instead by peripheral vasodilation (a reduction in systemic vascular resistance), suggesting preferential recruitment of efferent vagal vasomotor pathways over direct cardiac effects.
